# Hexokinase detachment from mitochondria drives the Warburg effect to support compartmentalized ATP production

**DOI:** 10.1101/2025.02.07.637120

**Authors:** Kimberly S. Huggler, Carlos A. Mellado Fritz, Kyle M. Flickinger, Gavin R. Chang, Meghan F. McGuire, Jason R. Cantor

## Abstract

Hexokinase (HK) catalyzes the synthesis of glucose-6-phosphate, marking the first committed step of glucose metabolism. Most cancer cells express two homologous isoforms (HK1 and HK2) that can each bind to the outer mitochondrial membrane (OMM). CRISPR screens across hundreds of cancer cell lines indicate that both are dispensable for cell growth in traditional culture media. By contrast, *HK2* deletion impairs cell growth in Human Plasma-Like Medium (HPLM). Here, we find that HK2 is required to maintain sufficient cytosolic (OMM-detached) HK activity under conditions that enhance HK1 binding to the OMM. Notably, OMM-detached rather than OMM-docked HK promotes “aerobic glycolysis” (Warburg effect), an enigmatic phenotype displayed by most proliferating cells. We show that several proposed theories for this phenotype cannot explain the *HK2* dependence and instead find that *HK2* deletion severely impairs glycolytic ATP production with little impact on total ATP yield for cells in HPLM. Our results reveal a basis for conditional *HK2* essentiality and suggest that demand for compartmentalized ATP synthesis underlies the Warburg effect.

---

The genetic dependencies of human cells are shaped by an interplay of intrinsic and extrinsic factors^1^. Defining such dependencies can generate insights into protein function and the molecular drivers of various cellular processes and also holds promise for nominating therapeutic targets^2–7^. CRISPR-based genetic screening across hundreds of cell lines has enabled the identification of core essential genes likely necessary for cell growth in most cases, while also revealing that other cancer vulnerabilities may vary with genotype or lineage^8–11^. However, despite mounting evidence that gene essentiality can depend on cell growth conditions as well^12–16^, there has been limited consideration into how the metabolic environment impacts the genetic dependencies of proliferating human cells. Moreover, most CRISPR screens are performed in vitro using traditional culture media that poorly resemble nutrient conditions that cells may encounter within the human body^17^.

We previously reported the development of Human Plasma-Like Medium (HPLM), a synthetic cell culture medium that contains over sixty metabolites and small ions at concentrations reflective of those found in human blood^18^. To prepare complete HPLM, we supplement the basal medium with 10% dialyzed fetal bovine serum (FBS) (HPLM^+dS^), which contributes various biomolecules that help to support cell growth while minimizing the addition of undefined polar metabolites. In a prior study, we tested the hypothesis that medium composition can affect the genetic dependencies of human cancer cells^19^. By performing CRISPR screens in HPLM^+dS^ and conventional media, we identified hundreds of conditionally essential candidate genes. For most cases, underlying causes that explain the conditional CRISPR phenotypes for such genes are not immediately apparent but could generate new insights into how specific proteins support cell growth and why such contributions vary based on nutrient conditions.

Hexokinase (HK) catalyzes the phosphorylation of glucose to glucose-6-phosphate (G6P), effectively trapping imported glucose inside the cell. This reaction marks the first committed step of glucose metabolism. Most cancer cells express two homologous isoforms of HK from distinct genes: *HK1* and *HK2*. Our previous CRISPR screen results revealed that hexokinase 2 *(HK2)* was selectively essential for cells in HPLM^+dS^ versus traditional media, but that *HK1* was dispensable in each case. Here, we uncover the basis for this selective dependence. By leveraging conditional *HK2* essentiality, follow-up work reveals a distinct rationale for why proliferating cells engage in aerobic glycolysis, a metabolic phenotype whose role in supporting cell growth has been extensively studied and debated for decades^20–24^. Our results establish further evidence that conditional essentiality in HPLM can be exploited to gain insights into protein function and cell metabolism, suggesting several directions for future study.

## RESULTS

### Conditional *HK2* essentiality can vary with intrinsic factors

We previously used a genome-wide single guide (sg)RNA library to perform CRISPR screens in K562 chronic myeloid leukemia (CML) cells grown in either HPLM^+dS^ or RPMI prepared with HPLM-defined glucose (5 mM) and 10% dialyzed FBS (RPMI^+dS^)^19^. Analysis of these screen results revealed that *HK2* was among the top 1% of hit genes that scored as selectively essential in HPLM^+dS^ (Fig. 1a). Notably, *HK2* was defined as a dependency in only 4% of the 1,100 CRISPR screens from DepMap^10^ (Extended Data Fig. 1a). *HK2* encodes one of five human enzymes that can catalyze the conversion of glucose to G6P but vary in normal tissue distribution, affinity for glucose, subcellular localization, number of catalytic domains, and relative sensitivity to product inhibition^25,26^. Upon HK-mediated transfer of a phosphate group from adenosine triphosphate (ATP) to glucose, G6P can enter multiple branches of glucose metabolism (Fig. 1b).

**Fig. 1.**
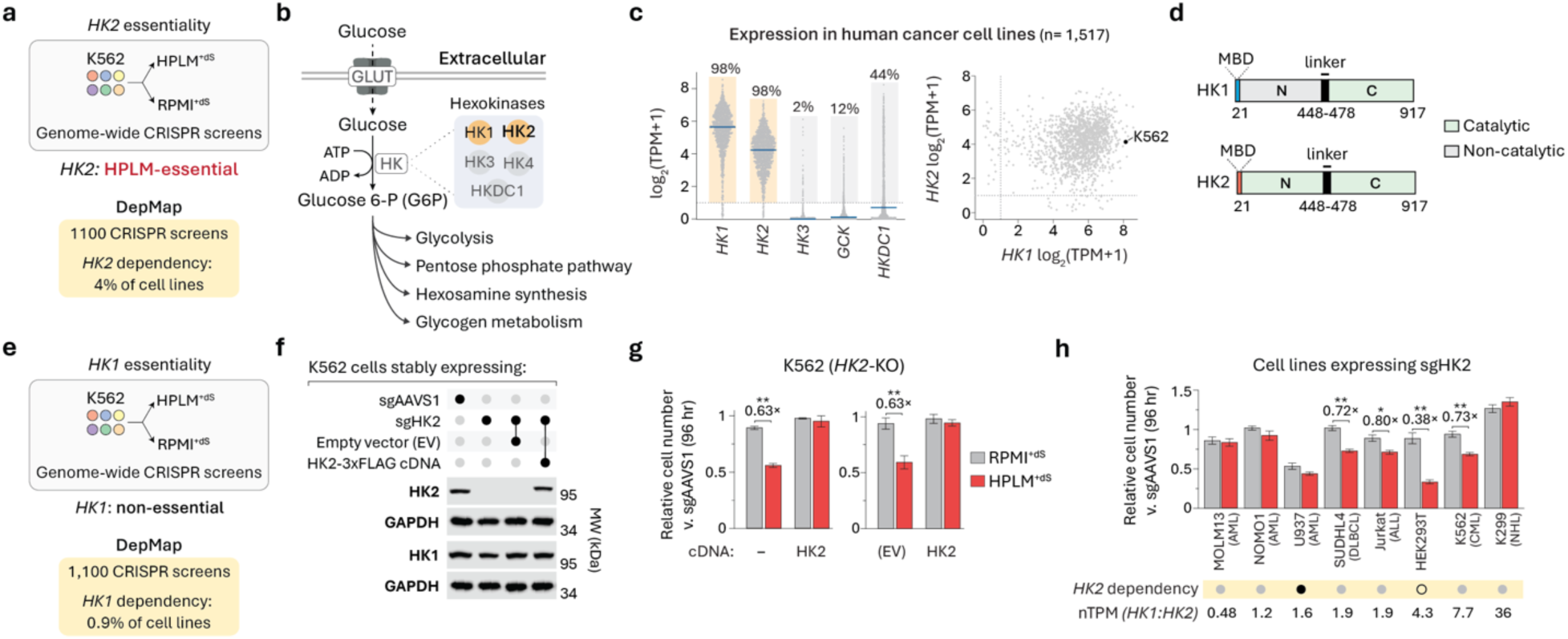
Conditional *HK2* essentiality can vary with intrinsic factors. (a and e) Dependency phenotypes for *HK2* (A) and *HK1* (e) from conditional essentiality profiling in K562 cells (top) and the DepMap project (bottom)^10^. (b) Schematic of the reaction catalyzed by hexokinase (HK) and possible metabolic fates of glucose 6-phosphate (G6P). Human HK isoforms that are expressed (orange) or absent (gray) in K562 cells. (c) Distribution of RNA transcript levels for each HK isoform across human cancer cell lines from cataloged RNA-Seq data^27^. Percentage, cell lines with log_2_(TPM+1) greater than 1. Blue line, median. (left) Comparison between *HK1* and *HK2* RNA transcript levels (right). TPM, transcripts per million. (d) Schematic of domain architecture for HK1 (top) and HK2 (bottom). Only the C-terminal domain of HK1 can phosphorylate glucose. MBD, mitochondrial binding domain. (f) Immunoblots for expression of HK1 and HK2 in *HK2*-knockout and control (sgAAVS1) K562 cells. GAPDH served as the loading control in both cases. (g) Growth of *HK2*-knockout versus control cells (mean ± SD, *n* = 3, \*\**P* < 0.005). EV, empty vector. (h) Relative growth of cells transduced with sgHK2 versus sgAAVS1 (mean ± SD, *n* = 3, \*\**P* < 0.005). Middle row, DepMap annotation for *HK2* dependency. Essential (black dot), non-essential (gray dot). HEK293T is not in the DepMap panel (unfilled). Bottom row, ratio of *HK1*-to-*HK2* RNA transcript levels from analyzed RNA-seq data^32^. nTPM, normalized TPM. ALL, acute lymphoblastic leukemia. AML, acute myeloid leukemia. CML, chronic myeloid leukemia. DLBCL, diffuse large B cell lymphoma. NHL, T cell non-Hodgkin’s lymphoma.

Cataloged RNA sequencing (RNA-seq) data indicate that *HK1* and *HK2* are almost uniformly co-expressed across a panel of over 1,500 cancer cell lines, whereas transcript levels for *HK3*, *GCK*, and *HKDC1* are negligible in most cases, including in K562 cells (Fig. 1c)^27^. Indeed, our screen results indicated that the latter three HK isoforms were dispensable for cell growth regardless of nutrient conditions – reflective of the general CRISPR phenotypes for these genes in DepMap (Extended Data Fig. 1b). In addition, we confirmed that HK1 and HK2 are both expressed in K562 cells and found that culture in HPLM^+dS^ versus RPMI-based media did not affect the total abundance of either protein (Extended Data Fig. 1c).

HK1 and HK2 share over 70% sequence identity and a comparable susceptibility to product (G6P) inhibition, and further, each displays sub-millimolar affinity for glucose^25^. Both proteins are also comprised of tandem domains joined by a linker helix: the C-terminal domain mediates glucose phosphorylation in each case, whereas the N-terminal domain is catalytic only in HK2^28,29^. Moreover, both isoforms contain an N-terminal 21-residue mitochondrial binding domain (MBD) that permits HK association with the outer mitochondrial membrane (OMM) (Fig. 1d)^30,31^. Despite the high level of structural and biochemical similarity between HK1 and HK2, our CRISPR screen results indicated that *HK1* was dispensable in both HPLM^+dS^ and RPMI^+dS^ – reflective of broadly cataloged data from the DepMap, where *HK1* was defined as essential for less than 1% of the profiled cell lines (Fig. 1e). Together, our results suggested that HK2 fulfills a non-redundant role for cell growth in HPLM^+dS^.

To begin to investigate the conditional CRISPR phenotype for *HK2*, we used an *HK2*-targeting sgRNA to engineer *HK2*-knockout K562 clonal cells, which showed HK1 levels comparable to control cells transduced with an *AAVS1*-targeting sgRNA (Fig. 1f). Consistent with our screen results, *HK2*-knockout caused a 40% stronger growth defect in HPLM^+dS^ versus RPMI^+dS^ (Fig. 1g). Importantly, the expression of an sgRNA-resistant *HK2* cDNA rescued this conditional impairment, while transducing these cells with an equivalent empty vector (EV) did not.

Since *HK2* has been rarely defined as a dependency in cataloged CRISPR screen results, we sought to determine whether the HPLM-essential phenotype of *HK2* may be conserved in additional cell lines. Using the two sgRNAs noted above, we transduced a panel of eight cell lines: K562, three additional blood cancer lines that we also previously screened in HPLM^+dS^ and RPMI^+dS^ but by using a focused sgRNA library^19^, one chosen to sample a distinct blood cancer type, one that shows nearly selective expression of *HK1* versus *HK2* according to analyzed RNA-seq data^32^, one that exhibits *HK2* dependence based on DepMap data, and one non-cancer adherent cell line (Extended Data Fig. 1d). When we evaluated the relative growth of these *HK2*-depleted cell lines, we found that half showed stronger growth defects in HPLM^+dS^ versus RPMI^+dS^, including K562 as expected (Fig. 1h).

Our focused sgRNA library screen results suggested that *HK2* was not conditionally essential in MOLM13 cells but scored as either a relatively weak (NOMO1) or strong (SUDHL4) HPLM-essential hit gene in the other two profiled cell lines. Consistent with these results, *HK2* depletion had similar conditional effects on the relative growth of K562 and SUDHL4 cells, but modest to negligible effects in the NOMO1 and MOLM13 lines. By contrast, *HK2* deletion caused a 50% growth defect in U937 cells regardless of culture in HPLM^+dS^ or RPMI^+dS^ but did not impair the relative growth of Karpas299 cells in either case – outcomes each perhaps as anticipated based on DepMap data (U937) or given the markedly higher transcript levels of *HK1* versus *HK2* (Karpas299). For the two remaining cell lines from our panel, *HK2* depletion led to stronger growth defects in HPLM^+dS^ versus RPMI^+dS^ – differential effects that were comparable (Jurkat) or even stronger (HEK293T) than those observed in K562 cells. Collectively, these results suggest that *HK2* essentiality can be shaped by an interplay of natural cell-intrinsic factors and nutrient conditions, wherein conditional *HK2* dependence was neither cell type-specific nor entirely correlated with the relative expression of *HK1* versus *HK2*.

### HK2 catalytic activity is necessary to support cell growth in HPLM

We next sought to determine a possible basis for conditional *HK2* essentiality by considering previous explanations for how HK may support cancer growth. HK1 and HK2 can each localize to the OMM in part via interaction with the voltage-dependent anion channel (VDAC)^33^. This association has been suggested to provide the OMM-bound HK with enhanced access to mitochondrial ATP released through a multi-protein channel that includes VDAC, enabling efficient glucose phosphorylation to specifically promote glycolysis^25,34–36^. However, this model has been questioned based on evidence suggesting that cytosolic ATP levels are far greater than the reported ATP affinities of HK1 or HK2^31,37^. In addition, prior work indicates that HK-VDAC binding may antagonize interactions between VDAC and pro-apoptotic proteins, effectively suppressing apoptosis^38–40^, though this role would unlikely be unique to HK2. Nonetheless, recent studies have reported that HK1 and HK2 can serve distinct non-catalytic roles as well^41,42^. Thus, we asked whether conditional *HK2* dependence might be linked to a non-catalytic function. To test this possibility, we engineered a putative kinase-dead *HK2* cDNA by mutating the catalytic aspartate residue in each HK2 domain (D209 and D657)^28^. Using a mass spectrometry (MS)-based assay to evaluate HK activity, we confirmed that our wild-type HK2 variant mediated time-dependent G6P synthesis, whereas our D209A-D657A variant showed no activity as expected (Fig. 2a and Extended Data Fig. 2a,b). When we transduced the kinase-dead *HK2* cDNA into our *HK2*-knockout cells, we observed no rescue of the relative growth defect in HPLM^+dS^ (Fig. 2b,c).

**Fig. 2.**
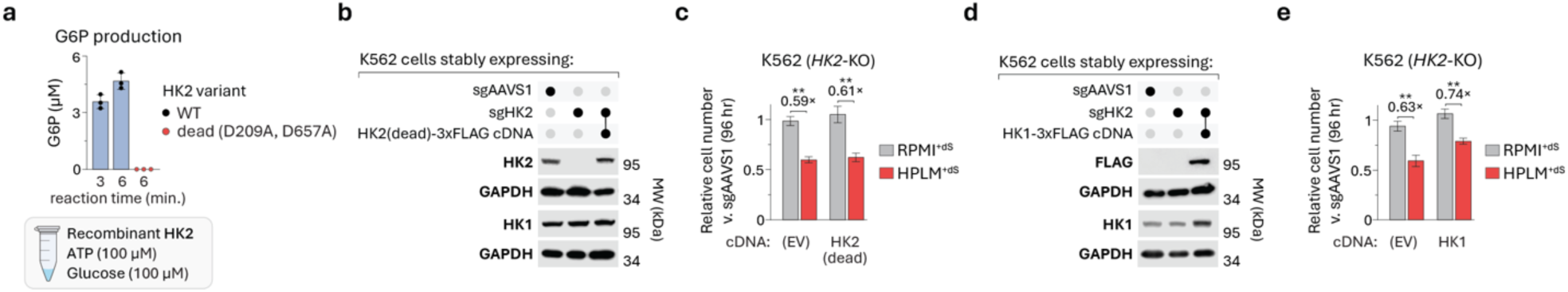
HK2 catalytic activity is necessary to support cell growth in HPLM. (a) Glucose 6-phosphate (G6P) levels measured from reactions of recombinant HK2 (either WT or the D209A, D657A variant) with glucose and ATP (mean ± SD, *n* = 3). WT, wild-type. (b and d) Immunoblots for expression of HK1 and HK2 (B) or HK1 and Flag (D) in *HK2*-knockout and control (sgAAVS1) K562 cells. GAPDH served as the loading control in all cases. (c and e) Relative growth of *HK2*-knockout versus control cells (mean ± SD, *n* = 3, \*\**P* < 0.005). EV, empty vector. HK2 (dead), D209A, D657A variant.

Next, we considered whether conditional *HK2* essentiality might be attributed to a limitation in total levels of high-affinity HK. The reported substrate affinities displayed by HK1 are comparable (ATP) or greater (glucose) versus those by HK2, and further, *K*_i_ values for G6P are similar between the two enzymes^25,28,43^. Thus, we sought to test if enforced overexpression of *HK1* could complement for loss of *HK2*. However, transducing an *HK1* cDNA into our *HK2*-knockout cells led to only a modest 10% rescue of the relative growth defect despite a nearly twofold increase in cellular HK1 levels (Fig. 2d, e). Collectively, these results suggested that mediating glucose phosphorylation is necessary but not sufficient to explain why *HK2* was differentially required for cell growth in HPLM^+dS^.

### Relative HK1-OMM binding varies with nutrient conditions

Prior work has suggested that the subcellular localization of HK shapes the metabolic fate of glucose, whereby OMM-docking promotes glycolysis but residing in the bulk cytosol diverts G6P to alternative branches of glucose metabolism such as the pentose phosphate pathway (PPP)^25,36,37,44^. However, these links have been derived in large part from (fusion) protein overexpression models or studies in cells that do not endogenously express both HK1 and HK2. Moreover, growing evidence indicates that HK-OMM binding is unlikely static and may instead be influenced by various factors^33^, including the states of different post-translational modifications on VDAC^45^, HK1^46^, or HK2^47^; glucose restriction^36^; intracellular pH^48^; and perhaps VDAC assembly within the OMM^49^. In addition, it is appreciated that protein overexpression can lead to its mislocalization inside cells^50^. Together, since excess HK1 partially compensated for loss of *HK2*, we hypothesized that relative HK-OMM binding might vary by isoform depending on culture in HPLM^+dS^ versus RPMI^+dS^.

To test this idea, we transduced wild-type K562 cells with either a *3xHA-eGFP-OMP25* (HA-MITO) or *3xMyc-eGFP-OMP25* (Control-MITO) cDNA^51^, and immunopurified mitochondria (mito-IP) from each respective population following growth in either HPLM^+dS^ or RPMI^+dS^. Immunoblot analysis confirmed selective enrichment of cytochrome c oxidase subunit 4 (COXIV) in the mito-IP from HA-MITO-expressing cells with only minimal detection of protein markers for the endoplasmic reticulum and peroxisomes – organelles that can directly interact with mitochondria^52–54^ (Fig. 3a). As expected, levels of HK1 versus HK2 in the mito-IP differed markedly despite the high degree of MBD sequence homology between these proteins, whereby HK2 was virtually undetectable regardless of culture in HPLM^+dS^ or RPMI^+dS^. Using COXIV as a mitochondrial control marker, we found that normalized HK1 levels in the mito-IP were sixfold higher than those of HK2 from the cells in HPLM^+dS^. Notably, similar analysis also revealed that mito-IP enrichment of HK1 was twofold lower from cells in RPMI^+dS^ versus HPLM^+dS^ with little difference in whole-cell HK1 levels otherwise, suggesting a shift in the subcellular distribution of HK1 favoring OMM detachment (Extended Data Fig. 3a, b). To determine whether *HK2* deletion might affect this conditional shift, we transduced the HA-MITO cDNA into our *HK2*-knockout cell lines harboring either *HK2* cDNA or EV, and then isolated mitochondria from each population. In both cases, we again observed a nearly twofold weaker enrichment of HK1 in the mito-IP from cells cultured in RPMI^+dS^ compared to HPLM^+dS^ (Fig. 3b, c). Together, these results indicate that nutrient conditions independent of defined glucose availability can influence HK1 translocation to the OMM regardless of HK2 expression.

**Fig. 3.**
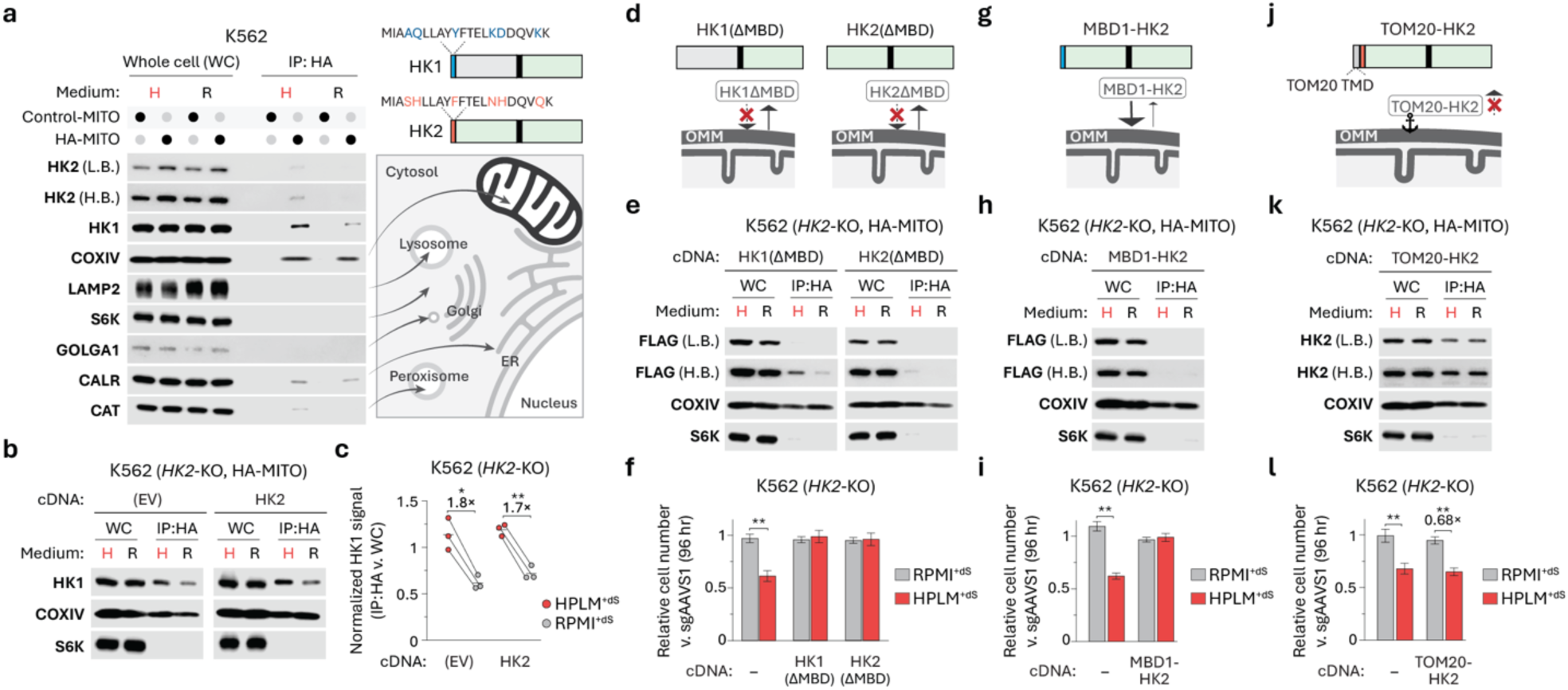
Relative HK1-OMM binding varies with nutrient conditions and HK detachment from mitochondria is necessary to support cell growth. (a) Immunoblots of indicated proteins in whole-cell (WC) lysates and mitochondria immunopurified by HA (IP:HA) from cells expressing 3xMyc-eGFP-OMP25 (Control-MITO) or 3xHA-eGFP-OMP25 (HA-MITO). L.B., low brightness. H.B., high brightness (left). HK1 and HK2 mitochondrial binding domain (MBD) sequences. Black text, positions of shared sequence identity (right, top). Schematic depicting putative subcellular localization of the indicated proteins (right, bottom). (b, e, h, and k) Immunoblots for HK1 (b), Flag signal (e and h), or HK2 (k) in WC lysates and IP:HA from *HK2*-knockout cells expressing HA-MITO. COXIV served as a mitochondrial control marker. S6K served as a non-mitochondrial control marker. EV, empty vector. MBD1, MBD sequence from HK1. (c) HK1 signal normalized by COXIV signal in IP:HA versus WC lysate based on immunoblot analysis (mean, *n* = 3, \*\**P* < 0.01, \**P* < 0.05). Each line connects points corresponding to the same biological replicate. Immunoblots in (b) depict a representative replicate. (d, g, and j) Schematic of domain architectures for MBD-deficient HK1 and MBD-deficient HK2 (d), MBD1-HK2 (g), and TOM20-HK2 (j). TOM20, Translocase of outer mitochondrial membrane 20. TMD, transmembrane domain (top). Anticipated influence of each modification on relative HK association with the outer mitochondrial membrane (OMM) (bottom). (f, i, and l) Relative growth of *HK2*-knockout versus control cells (mean ± SD, *n* = 3, \*\**P* < 0.005).

### HK detachment from mitochondria is necessary to support cell growth

Since our data suggested that *HK2* deletion impairs cell growth under conditions that lead to greater HK1-OMM binding, we next hypothesized that conditional *HK2* essentiality might be linked to a limitation in activity specific to OMM-detached HK. To test this idea, we transduced either an *HK1* or *HK2* cDNA lacking the respective N-terminal MBD into *HK2*-knockout cells harboring the HA-MITO cDNA (Fig. 3d and Extended Data Fig. 3c). When we isolated mitochondria from these populations, we found that MBD removal did not fully abolish HK1 detection in the mito-IP but markedly reduced mito-IP enrichment of HK1 by eightfold in HPLM^+dS^ and by nearly thirtyfold in RPMI^+dS^ based on signal from the fused Flag tag (Fig. 3e and Extended Data Fig. 3d). Moreover, Flag signal for MBD-deficient HK2 in the mito-IP was fivefold lower versus that for the equivalent HK1 variant following growth in HPLM^+dS^ but was otherwise undetectable in isolated mitochondria from cells in RPMI^+dS^. Consistent with our rationale and these mito-IP results, we found that expression of either truncated *HK* cDNA fully restored the relative growth of our *HK2*-knockout cells in HPLM^+dS^ (Fig. 3f).

Next, we considered whether increasing HK2-OMM binding may diminish the contribution of HK2 for cell growth. Given the differential enrichment of HK1 versus HK2 docked at the OMM based on our mito-IP data, we transduced *HK2*-knockout cells harboring HA-MITO cDNA with an *HK2* cDNA in which we replaced the endogenous MBD with the respective sequence from HK1 (MBD1) (Fig. 3g). However, we could not detect Flag signal for this HK2 variant in a mito-IP from cells grown in HPLM^+dS^ or RPMI^+dS^ (Fig. 3h and Extended Data Fig. 3e). Consistent with this apparent yet unanticipated lack of OMM association, *HK2*-knockout cells expressing MBD1-HK2 did not display a conditional growth defect in HPLM^+dS^ (Fig. 3i). Together, these results suggest that MBD1 is critical though not sufficient for HK1-OMM binding, and also that the MBD1 sequence cannot act as a broad OMM ‘binding factor’ when attached to other proteins. Indeed, a previous study reported that fusing an extended version of this domain (N-terminal 34 residues of HK1) enabled OMM-binding of green fluorescent protein in rodent cells but failed to drive similar localization for human glucokinase^30^.

We then sought an approach to anchor HK2 at the OMM. Translocase of outer mitochondrial membrane 20 (TOM20) is part of the TOM complex that imports proteins into mitochondria^55^. TOM20 contains an N-terminal transmembrane domain for proper localization, which has also been used to attach orthogonal proteins to the OMM^56,57^. Thus, we transduced *HK2*-knockout cells harboring the HA-MITO cDNA with a cDNA in which we fused the N-terminal encoding sequence of TOM20 to *HK2* (Fig. 3j). As expected, the TOM20-HK2 variant was markedly enriched in mitochondria isolated from these cells, with detection levels largely unaffected by culture in HPLM^+dS^ versus RPMI^+dS^ (Fig. 3k and Extended Data Fig. 3f). Moreover, expression of our *TOM20-HK2* cDNA had negligible impact on the growth impairment of *HK2*-knockout cells in HPLM^+dS^ (Fig. 3l). Of note, using our MS-based assay for HK activity, we also established that our TOM20-HK2 variant could phosphorylate glucose, further suggesting that this lack of rescue could likely be attributed to a shift in HK2 localization rather than to an unintended loss of catalytic activity (Extended Data Fig. 3g, h).

We also asked whether increasing glucose availability might compensate for the loss of *HK2* by promoting enhanced HK activity despite a seemingly unfavorable subcellular distribution of HK1. Nonetheless, preparing HPLM with twofold more glucose (10 mM) did not alter the relative growth of *HK2*-knockout cells in HPLM^+dS^ (Extended Data Fig. 3i). Overall, these results suggest that total OMM-detached (“cytosolic”) HK activity, as mediated by the two high-affinity HK isoforms (HK1 and HK2) co-expressed in nearly all human cancer lines, must meet a critical threshold to support cell growth.

### HK detachment from mitochondria supports aerobic glycolysis

Since our data indicated that HK2 is conditionally required to help drive “cytosolic” glucose phosphorylation, we next examined how loss of *HK2* affects glucose metabolism. Most proliferating cells display a tendency to ferment glucose even in the presence of oxygen, a phenotype known as “aerobic glycolysis” (or the Warburg effect in mammalian cells^58^) that is further defined by increased glucose consumption and elevated secretion of lactate (mammalian cells) or ethanol (microbes)^22,23^. Supporting this phenotype in cancer cells has been a role assigned to HK2 but often on the basis that translocation to the OMM is required^44,59–62^. Although we found that OMM-detached HK promotes cell growth, we sought to determine whether *HK2* deletion differentially affects glucose fermentation in HPLM^+dS^ versus RPMI^+dS^. To assess this, we collected conditioned media at several time points along growth curves and then measured specific growth rates and net rates of metabolite exchange across the log phase (Fig. 4a). Similar to the case for wild-type K562 cells^18^, the growth rate of our control cells was only modestly reduced in HPLM^+dS^ relative to RPMI^+dS^ (Extended Data Fig. 4a). In addition, *HK2*-knockout cells displayed a 20% slower growth rate specific to culture in HPLM^+dS^, a defect that could be rescued by the expression of different HK1/2 variants based on corresponding “cytosolic” activity (Extended Data Fig. 4b). Nonetheless, *HK2* deletion markedly impaired the rates of glucose uptake and lactate secretion by 35 to 40% during log growth in RPMI^+dS^ – effects exacerbated by an additional 10 to 15% in HPLM^+dS^ but reversed upon expression of our *HK2* cDNA (Fig. 4b, c). Moreover, these defects could be similarly restored by expressing other HK1/2 variants, again based on relative OMM detachment (Fig. 4d, e). Interestingly, we also found that the ratio of lactate release to glucose uptake was comparable in all cases, with values ranging from 1.3 to 1.5, suggesting that these cells secreted 65 to 75% of incoming glucose as lactate regardless of *HK2* expression, relative cytosolic HK activity, or culture in HPLM^+dS^ versus RPMI^+dS^ (Fig. 4f and Extended Data Fig. 4c).

**Fig. 4.**
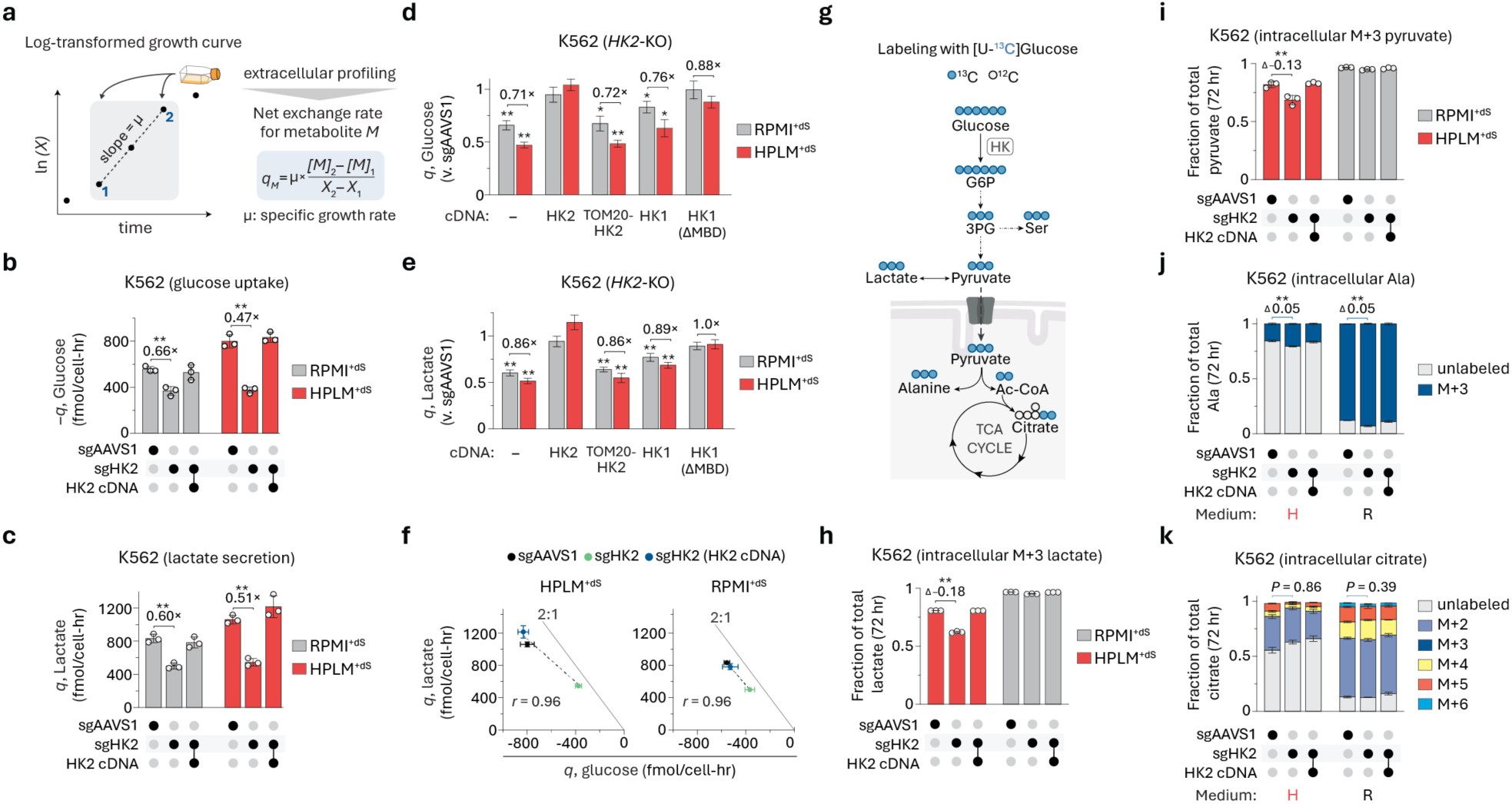
HK detachment from mitochondria supports aerobic glycolysis. (a) Schematic depicting a log-transformed growth curve. Gray box highlights points during log phase, whereby point 1 was 24 hr and point 2 was 72 hr for all experiments. *X*, cell number. μ, specific growth rate (left). Equation used to calculate the rate of net exchange *(q)* for a given extracellular metabolite *(M)* between two time points (right)^122^. (b and c) Absolute rates of net exchange for glucose (b) and lactate (c) in *HK2*-knockout and control cells (mean ± SD, *n* = 3, \*\**P* < 0.005). Negative values, net uptake. Positive values, net secretion. (d and e) Relative net exchange for glucose (d) and lactate (e) in *HK2*-knockout versus control cells (mean ± SEM, *n* = 3, \*\**P* < 0.005, \**P* < 0.05). (f) Comparison of net exchange rates for glucose versus lactate in *HK2*-knockout and control cells (mean ± SEM, *n* = 3). Gray line denotes the theoretical maximum of two lactate molecules released per glucose molecule imported (2:1). *r*, Pearson’s correlation coefficient. (g) Schematic depicting the incorporation of ^13^C from [U-^13^C] glucose into various metabolites. G6P, glucose 6-phosphate. 3PG, 3-phosphoglycerate. (h-k) Fractional labeling of lactate (h) pyruvate (i), alanine (j), and citrate (k) in *HK2*-knockout and control cells (mean ± SD, *n* = 3, \*\**P* < 0.005). Values above brackets indicate differences in fractional labeling between bars (h-j). Comparisons are for M+2-citrate (K). M+X, incorporation of X ^13^C.

To determine whether loss of *HK2* alters glucose utilization, we evaluated the ^13^C labeling of various metabolites following cell growth in HPLM^+dS^ or RPMI^+dS^ prepared with [U-^13^C] glucose (Fig. 4g). Notably, *HK2-*knockout decreased the fraction of lactate labeled with three ^13^C (M+3) by 20% for cells grown in HPLM^+dS^, supporting rationale that HK2 promotes glucose fermentation to lactate (Fig. 4h). By contrast, loss of *HK2* did not affect the nearly complete M+3-labeling of lactate for cells grown in RPMI^+dS^, indicating that glucose was the lone source of this pool as expected given the distinct lack of defined lactate provided by RPMI versus HPLM. Moreover, the M+3-labeling patterns for pyruvate largely resembled those found for lactate, whereby *HK2* deletion led to a similar but slightly smaller reduction in M+3-pyruvate for cells in HPLM^+dS^ but had little effect on this labeling for cells in RPMI^+dS^, which also distinctly lacks defined pyruvate (Fig. 4i). These results suggested that cytosolic pyruvate and lactate were in equilibrium, with the slight difference in M+3-labeling (5%) likely attributed to an added contribution of mitochondrial pyruvate derived from a non-glucose source. This analysis also revealed that *HK2* deletion caused modest yet comparable effects on glucose diversion into serine synthesis regardless of growth in HPLM^+dS^ or RPMI^+dS^, with M+3-serine labeling boosted by 6% in both cases (Extended Data Fig. 4d). We also examined other products of pyruvate metabolism beyond lactate. Control cells grown in RPMI^+dS^ displayed markedly higher M+3-alanine labeling as expected given that HPLM uniquely contains defined alanine (430 μM), but *HK2*-knockout slightly elevated this fractional labeling by 5% in each condition (Fig. 4j). Moreover, total ^13^C labeling of citrate was much greater from control cells grown in RPMI^+dS^ versus HPLM^+dS^, including for the M+2 fraction often used to characterize glucose entry into the tricarboxylic acid (TCA) cycle, though loss of *HK2* had minimal effects on the fractional labeling of citrate in either condition (Fig. 4k).

We also asked whether *HK2* deletion affects diversion of G6P into the oxidative branch of the PPP (ox-PPP) by assessing ^13^C patterns for 3-phosphoglycerate (3PG) following growth in HPLM^+dS^ or RPMI^+dS^ prepared with [1,2-^13^C] glucose (Extended Data Fig. 4e). Although prior work suggested that HK detachment from the OMM promotes this G6P diversion^36,37^, our analysis revealed that 3PG pools were comprised of nearly equivalent unlabeled and M+2 fractions in all cases, indicating that ox-PPP activity was negligible regardless of either *HK2* expression or nutrient conditions (Extended Data Fig. 4f)^63^. By contrast, *HK2* deletion reduced the M+2 fractional labeling of pyruvate and lactate specific to cells grown in HPLM^+dS^ as expected based on our [U-^13^C] glucose tracing data (Extended Data Fig. 4g, h). Collectively, these results demonstrated that OMM-detached rather than OMM-docked HK correlates with defining features of the Warburg effect and that HK2 promoted glucose fermentation but had limited effects on glucose utilization otherwise.

### Conditional *HK2* essentiality cannot be traced to a direct gene-nutrient interaction

Next, we sought to identify the difference in composition between HPLM and RPMI that may underlie conditional *HK2* dependence. Interestingly, although we normalized glucose availability in RPMI to match HPLM, our data indicated that rates of glucose uptake displayed by control cells were 40% higher in HPLM^+dS^ (Fig. 5a). Nonetheless, exogenous glucose levels measured during log growth were comparable between the two conditions, perhaps suggesting cellular preference or differential reliance on alternate substrates in RPMI^+dS^ (Extended Data Fig. 5a). Glutamine is the most abundant amino acid in blood and serves not only as a critical nitrogen donor for macromolecule biosynthesis, but also as an important carbon source upon its two-step conversion to α-ketoglutarate (α-KG) (Fig. 5b)^64,65^. Interestingly, when we measured rates of net exchange for glutamine using our growth curve data, we found that baseline uptake rates were 50% higher in RPMI^+dS^ and also slightly increased with *HK2*-knockout specific to RPMI^+dS^ (Fig. 5c, d). Notably, RPMI contains fourfold more glutamine than HPLM. Thus, we hypothesized that this difference might cause cells to display a stronger reliance on glucose in HPLM^+dS^ versus RPMI^+dS^, and in turn, a greater dependence on *HK2*. However, normalizing the glutamine levels between HPLM and RPMI had no effect on the relative growth of *HK2*-knockout cells regardless of which basal medium was modified (Fig. 5e).

**Fig. 5.**
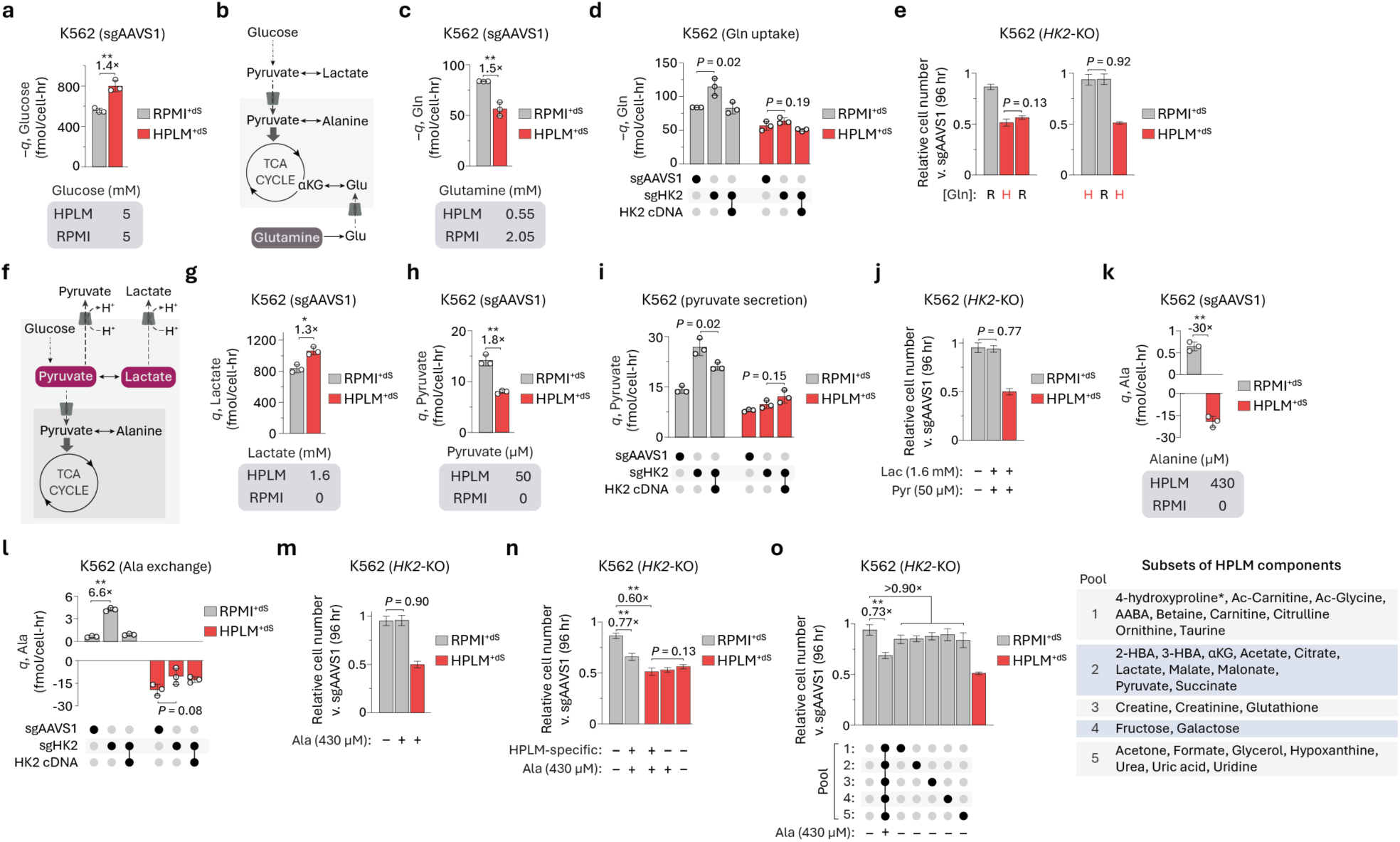
Conditional *HK2* essentiality cannot be traced to a direct gene-nutrient interaction. (a, c, g, h, and k) Absolute rates of net exchange for glucose (a), glutamine (c), lactate (g), pyruvate (h), and alanine (k) in control cells (mean ± SD, *n* = 3, \*\**P* < 0.005, \**P* < 0.01) (top). Defined levels of each respective metabolite in HPLM and RPMI (bottom). (b) Schematic depicting glutamine entry into the TCA cycle as α-ketoglutarate (α-KG). (d, i, and l) Absolute rates of net exchange for glutamine (d), pyruvate (i), and alanine (l) in *HK2*-knockout and control cells (mean ± SD, *n* = 3, \*\**P* < 0.005). (e, j, m, and n) Relative growth of *HK2*-knockout versus control cells (mean ± SD, *n* = 3, \*\**P* < 0.005). H, HPLM-defined concentrations; R, RPMI-defined concentrations (e). (f) Schematic depicting the proton-coupled secretion of pyruvate and lactate. (o) Relative growth of *HK2*-knockout versus control cells (mean ± SD, *n* = 3, \*\**P* < 0.005) (left). Pools of defined HPLM components. *RPMI contains defined 4-hydroxyproline at a concentration (153 μM) nearly eightfold higher than in HPLM. See Supplementary Table 1 (right).

We then considered the availability of metabolites related to the other defining characteristic of aerobic glycolysis (Fig. 5f). Previous work suggests that increased lactate secretion can help cells maintain intracellular pH and redox balance for growth^66–69^. Similar to the case for glucose uptake, we found that lactate secretion rates for control cells were 30% higher in HPLM^+dS^ despite the distinct inclusion of lactate (1.6 mM) in HPLM relative to RPMI (Fig. 5g and Extended Data Fig. 5b). In addition, pyruvate release may enable cells to remove excess protons generated from glycolysis as well, and notably, prior work found that blocking this export could impair cell growth^70^. While pyruvate is also only defined in HPLM (50 μM) versus RPMI, our net exchange analysis revealed that baseline pyruvate secretion rates were twofold higher in RPMI^+dS^ versus HPLM^+dS^ and also modestly increased by *HK2*-knockout, though this added effect was only partially reversed with the expression of our *HK2* cDNA (Fig. 5h, i). Together, we reasoned that differences in the defined levels of pyruvate or lactate might exacerbate the effects of *HK2* deletion by altering concentration gradients necessary to sustain pH homeostasis. However, the combined addition of both metabolites at HPLM-defined levels had little impact on the relative growth of *HK2*-knockout cells in RPMI^+dS^ (Fig. 5j).

Pyruvate can also be reversibly converted to alanine in a reaction localized to mitochondria in most cell lines^19^. Beyond marked disparities in relative glucose contributions to alanine synthesis, we found that control cells grown in RPMI^+dS^ secreted alanine, whereas those in HPLM^+dS^ consumed alanine and at far higher rates (Fig. 5k). Moreover, *HK2* deletion had little impact on alanine uptake in HPLM^+dS^ but boosted the alanine secretion rates in RPMI^+dS^ by nearly sevenfold – an effect reversed by the expression of different HK1/2 variants again based on respective cytosolic activity (Fig. 5l and Extended Data Fig. 5c). This suggested that *HK2*-knockout cells could perhaps be better poised to eliminate excess pyruvate as alanine if fermentation is impaired. Nonetheless, supplementing RPMI with HPLM-defined alanine did not affect the relative growth of *HK2*-knockout cells (Fig. 5m).

Since our rationale-driven approaches did not uncover a possible gene-nutrient interaction, we then tested HPLM derivatives with full sets of either amino acids or salts adjusted to match RPMI-defined levels. However, these unbiased modifications similarly failed to rescue the relative growth defect of our *HK2*-knockout cells (Extended Data Fig. 5d). HPLM also contains over 30 metabolites not otherwise defined in RPMI, including pyruvate, lactate, and alanine (within the categorized set of amino acids). Notably, removing alanine in isolation or in combination with all other HPLM-specific metabolites had no impact on the relative growth of our *HK2*-knockout cells in HPLM^+dS^, whereas the combined addition of these components to RPMI caused a partial impairment equivalent to 60% of the growth defect in HPLM^+dS^ versus RPMI^+dS^ (Fig. 5n). Surprisingly, however, when we tested various RPMI derivatives containing subsets of these HPLM-specific components, we found that none could recapitulate the effect induced upon their combined addition, as only slight and comparable growth defects were observed in each case (Fig. 5o).

Collectively, these results suggest that conditional *HK2* essentiality is shaped by a complex and combinatorial set of differences in medium composition that might in part alter HK1 binding to the OMM, whereby even the combined addition of HPLM-specific metabolites led to distinct effects on the growth of *HK2*-knockout cells depending on which basal medium was supplemented. Of note, while we prepared HPLM from manually constructed stock solutions for all experiments, we found that the conditional growth phenotype for *HK2*-knockout K562 cells was preserved and comparable when we instead prepared HPLM^+dS^ by using commercial HPLM (Thermo) (Extended Data Fig. 5e).

### HK detachment from mitochondria drives glycolytic ATP production

Since our data revealed that HK detachment from the OMM promotes aerobic glycolysis, we next reasoned that conditional *HK2* essentiality could serve as a node to investigate why the Warburg effect is beneficial for cell growth. Among several such proposed theories is that proliferating cells use rapidly consumed glucose as a carbon source to support the biosynthesis of macromolecules such as lipids, nucleotides, and hexosamines (Extended Data Fig. 6a)^20,21^. Thus, although our results suggested that K562 cells ultimately secrete up to 75% of imported glucose as lactate regardless of *HK2* expression or nutrient conditions, we considered whether the loss of *HK2* might limit biomass production. To test this idea, we generated HPLM derivatives intended to compensate for impaired utilization of glucose as a carbon source by feeding alternative biosynthetic pathways or providing critical pathway intermediates (Fig. 6a). N-acetylglucosamine can be used as a salvage substrate for hexosamine biosynthesis. Hypoxanthine can support the purine salvage pathway and is otherwise a defined HPLM component (10 μM), while adenosine and guanosine are nucleosides that may each act as substrates in this pathway as well. Uridine is another HPLM-specific component (3 μM) and, like cytidine, a nucleoside that can enter the pyrimidine salvage pathway. Thymidine is a nucleoside that can be phosphorylated to form thymidylate, a precursor of deoxythymidine triphosphate – a key substrate for DNA synthesis. Acetate may be used as another carbon source for lipid synthesis and, like hypoxanthine and uridine, is also a defined component of HPLM (40 μM). Dimethyl α-KG is a cell-permeable form of α-KG that could perhaps directly provide carbons to the TCA cycle. Overall, when we evaluated the relative growth of our *HK2*-knockout cells in HPLM derivatives containing subsets of these metabolites at supraphysiologic levels, we observed minimal effects in all cases (Fig. 6b). These results suggested that the critical role of OMM-detached HK was unlikely linked to supporting macromolecule synthesis, consistent with rationale that excreting most imported glucose as lactate does not efficiently provide carbon for anabolic pathways^22^.

**Fig. 6.**
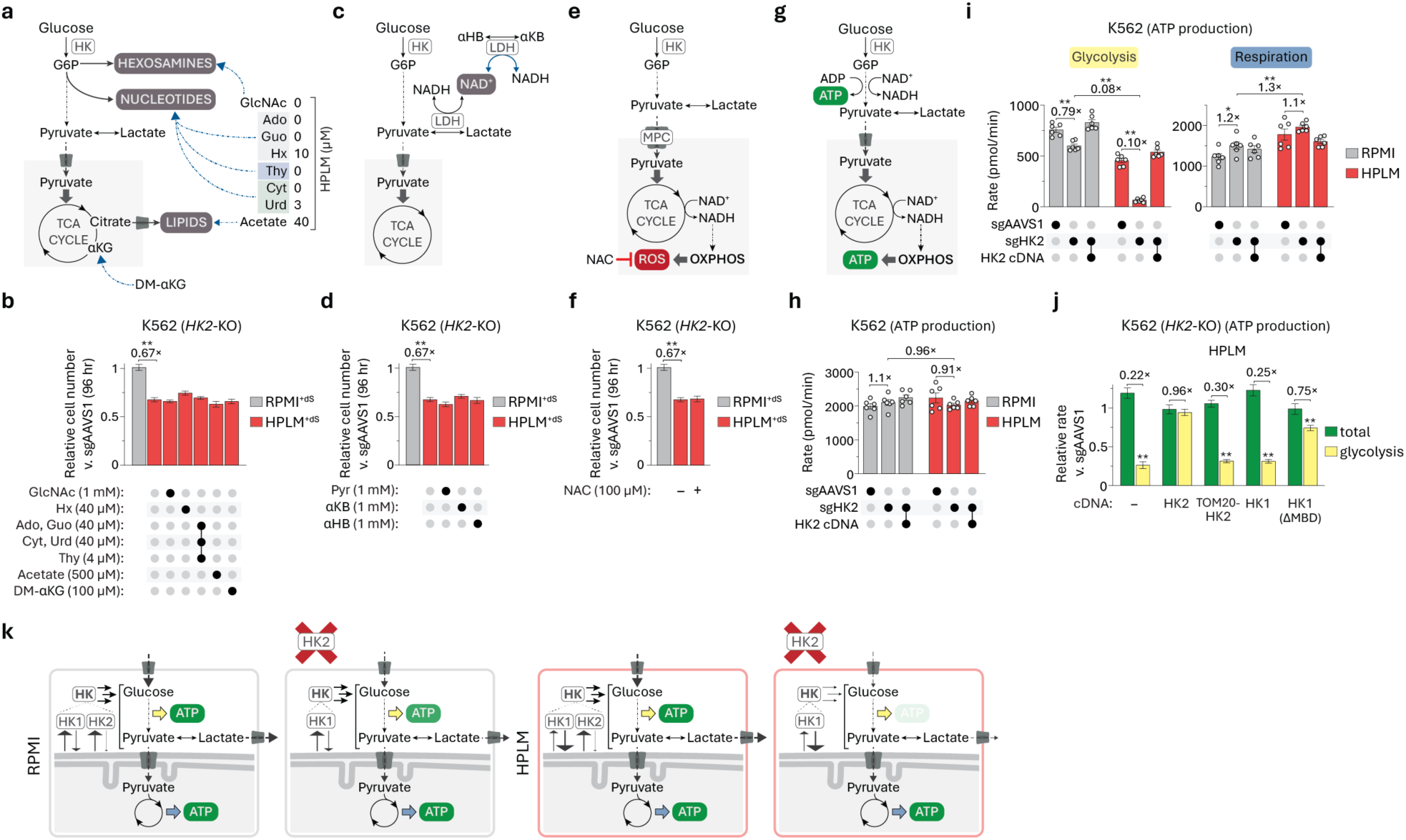
HK detachment from mitochondria drives glycolytic ATP production. (a) Glucose can be used as a carbon source for the biosynthesis of macromolecules such as lipids, nucleotides, and hexosamines. Schematic depicting various metabolites that can serve as salvage substrates or otherwise provide key intermediates to support these biosynthetic pathways. HPLM-defined concentrations of each metabolite are displayed. α-KG is also a defined HPLM component. GlcNAc, N-acetylglucosamine. Ado, adenosine. Guo, guanosine. Hx, hypoxanthine. Thy, thymidine. Cyt, cytidine. Urd, uridine. DM-α-KG, dimethyl α-KG. (b, d, and f) Relative growth of *HK2*-knockout versus control cells (mean ± SD, *n* = 3, \*\**P* < 0.005). Pyr, pyruvate. αKB, α-ketobutyrate. αHB, α-hydroxybutyrate (d). NAC, N-acetylcysteine (f). (c) Pyruvate and αKB can each serve as an electron acceptor for the reaction catalyzed by lactate dehydrogenase (LDH). αHB (or 2-hydroxybutyrate), which is defined in HPLM at 50 μM, can act as an electron donor for this reaction. (e) Reactive oxygen species (ROS) are byproducts of oxidative phosphorylation (OXPHOS). NAC is a potent antioxidant that can scavenge ROS. (g) Glucose metabolism can generate ATP by glycolysis or by respiration (OXPHOS). (h) Total ATP production rates by glycolysis and respiration combined in *HK2*-knockout and control cells (mean ± SEM, *n* = 6). HPLM and RPMI used for ATP production assays were not supplemented with dialyzed fetal bovine serum. See Supplementary Table 1. (i) Rates of ATP production by glycolysis (left) and by respiration (right) in *HK2*-knockout and control cells (mean ± SEM, *n* = 6, \*\**P* < 0.005, \**P* < 0.05). (j) Relative rates of ATP production by glycolysis (yellow) or by respiration and glycolysis combined (green) in *HK2*-knockout versus control cells (mean ± SEM, *n* = 6, \*\**P* < 0.005). (k) Proposed model of conditional *HK2* essentiality based on differential cytosolic HK activity driven by relative HK1-OMM binding in RPMI versus HPLM.

Prior work based on regulating the activity of pyruvate dehydrogenase suggested that aerobic glycolysis allows proliferating cells to support a limiting demand for NAD^+^ regeneration from NADH^71^. Nonetheless, we found that the pyruvate-to-lactate ratio – a proxy for the cytosolic NAD^+^-to-NADH ratio^68^ – was twofold higher in *HK2*-knockout versus control cells regardless of growth in HPLM^+dS^ or RPMI^+dS^ (Extended Data Fig. 6b, c). In addition, providing supraphysiologic levels of either an electron acceptor (pyruvate; α-ketobutyrate) or donor (α-hydroxybutyrate) for lactate dehydrogenase had little impact on the relative growth of *HK2*-knockout cells in HPLM^+dS^ (Fig. 6c, d). Moreover, treating these cells with duroquinone, which can serve as an electron acceptor for a distinct NADH-oxidizing reaction^71^, also failed to provide a rescue effect and instead elicited comparable growth defects in each condition (Extended Data Fig. 6d).

Another proposed theory for the Warburg effect is that glucose fermentation allows cells to limit the production of reactive oxygen species – byproducts of mitochondrial metabolism whose buildup can lead to oxidative damage and impair cell growth^72–74^. To test this rationale as a basis for conditional *HK2* dependence, we supplemented HPLM with a potent antioxidant (N-acetylcysteine), which similarly failed to restore the relative growth of our *HK2*-knockout cells (Fig. 6e, f). Additionally, previous studies reported that impeding the mitochondrial import of pyruvate could promote aerobic glycolysis^75,76^, whereas others suggested that inhibiting the mitochondrial pyruvate carrier (MPC) had little impact on either glucose uptake or lactate secretion^77,78^. In prior work, we identified both MPC components *(MPC1* and *MPC2)* as conditionally essential for cells grown in RPMI^+dS^ versus HPLM^+dS^ – phenotypes traced to differential alanine availability^19^. When we tested an MPC inhibitor (UK5099) against our control cells, we indeed observed a 50% growth defect specific to treatment in RPMI^+dS^ as expected (Extended Data Fig. 6e, f). Nonetheless, UK5099 treatment failed to improve the growth of *HK2*-knockout cells in HPLM^+dS^, though *HK2* deletion partially alleviated UK50999 sensitivity in RPMI^+dS^ perhaps on the basis that reduced fermentation can divert more pyruvate into mitochondria, leading to a boost in de novo alanine synthesis.

Although respiration generates over tenfold more ATP per glucose molecule than glycolysis, other explanations for the Warburg effect have been guided by models suggesting that glycolytic ATP production is either the faster or more proteome efficient process^79–83^. However, these theories have also been debated, in part on rationale that proliferating cells may not be ATP limited and also given recent work indicating that respiration is the more proteome efficient pathway for ATP synthesis^23,84^. Consistent with this, we found that *HK2* deletion had no effect on cellular ATP levels following growth in either HPLM^+dS^ or RPMI^+dS^ (Extended Data Fig. 6g). Nonetheless, we reasoned that considering ATP yield alone might mask the possibility that loss of *HK2* differentially impacts relative ATP production by glycolysis or respiration. Thus, we delineated the rates of ATP synthesis from each pathway (Fig. 6g). Our analysis revealed that total ATP production rates were largely unaffected by *HK2* expression or culture in HPLM versus RPMI but that, indeed, the relative pathway contributions varied (Fig. 6h,i). First, we found that baseline rates of ATP synthesis from respiration were faster than by glycolysis in both RPMI (1.6-fold) and HPLM (4-fold), and further, that baseline rates of glycolytic ATP production were 1.7-fold higher in RPMI versus HPLM (Extended Data Fig. 6h). Interestingly, *HK2* deletion shifted relative pathway contributions to ATP synthesis by roughly 20% favoring respiration for cells in RPMI, but severely impaired glycolytic ATP production rates (10-fold) coupled with only a modest boost in mitochondrial ATP generation for cells in HPLM – effects reversed with expression of our *HK2* cDNA. Together, glycolytic ATP synthesis rates were 10-fold lower for *HK2*-knockout cells in HPLM versus RPMI. Notably, this marked defect could also be rescued by the expression of distinct HK1/2 variants again based on respective cytosolic HK activity (Fig. 6j).

Collectively, our results suggest a model in which HK detachment from the OMM promotes aerobic glycolysis (Warburg effect) for compartmentalized ATP production. *HK2* deletion leads to a reduction in the “cytosolic” HK activity that helps to drive this metabolic phenotype. However, such effects are exacerbated in HPLM^+dS^ versus RPMI^+dS^ based on conditional differences in relative HK1 docking at the OMM. In turn, rates of glycolytic ATP production fall below a critical threshold needed to sufficiently support growth in HPLM^+dS^ despite minimal impacts on bulk ATP levels and total ATP synthesis rates (Fig. 6k). Ultimately, this localized defect cannot be compensated by the diffusion of mitochondrial-derived ATP in the cytosol, leading to an impairment of at least one energy-demanding essential processes that relies on the proximal source of ATP generated by glycolysis.

### 2-deoxyglucose efficacy correlates with HK detachment from mitochondria

2-deoxyglucose (2-DG) is a synthetic glucose analog that has been tested in clinical trials for cancer therapy^85^. Beyond competing with glucose for HK activity, 2-DG can be phosphorylated by HK to produce 2-deoxyglucose-6-phosphate (2-DG6P), which cannot be further metabolized inside the cell and thus accumulates. Acting as a G6P analog, 2-DG6P competitively disrupts G6P isomerase activity and also allosterically inhibits HK (Extended Data Fig. 7a)^86^. Although the clinical success of 2-DG has been limited, as largely attributed to its small therapeutic window, we asked whether *HK2* deletion might differentially affect 2-DG sensitivity in cells that otherwise co-express HK1. When we tested 2-DG against our control K562 cells at a single dose, we observed comparable 50% growth defects in both HPLM^+dS^ and RPMI^+dS^ as expected given that 2-DG is considered a non-selective HK inhibitor (Extended Data Fig. 7b). However, *HK2*-knockout effectively reduced the relative response to 2-DG by 25% only in HPLM^+dS^. These results suggest that 2-DG may be more efficiently captured by HK1/2 residing within the (bulk) cytosol than docked at the OMM regardless of the HK isoform.

## DISCUSSION

### Conditional HK2 essentiality reveals dependence on HK detached from mitochondria

We previously identified *HK2* as a selectively essential gene for K562 leukemia cells in HPLM versus RPMI with equivalent glucose availability^19^. Here, our study reveals that HK2 is required for “cytosolic” glucose phosphorylation under conditions where this spatially dependent activity would become limiting based on relative HK1 binding to the OMM. Previous studies have proposed that this HK-OMM interaction promotes glycolysis, while HK detachment from the OMM instead directs G6P to other pathways^25,36,37,44^. However, we find that OMM-detached (“cytosolic”) HK activity correlates with glucose uptake and lactate secretion – defining characteristics of aerobic glycolysis (Warburg effect) – and glycolytic ATP production. By contrast, *HK2* deletion has little impact on ATP yield, total ATP synthesis rates, and non-glycolytic branches of glucose metabolism. Notably, we find that only a negligible fraction of HK2 is associated with mitochondria from cells grown in either HPLM or RPMI, whereas mitochondrial enrichment of HK1 is twofold lower in RPMI versus HPLM regardless of *HK2* expression. Our results suggest that conditional *HK2* essentiality can be attributed to this differential enrichment of HK1 at the OMM, but further work is necessary to determine a molecular basis for how nutrient conditions influence the HK1-OMM interaction.

We also found that *HK2* depletion did not differentially impair growth in HPLM versus RPMI in all cases across an extended cell line panel, though these effects were neither cell type-specific nor dictated by reported expression levels of *HK1* versus *HK2*. In addition, we could not trace conditional *HK2* dependence to the differential availability of just a single component between HPLM and RPMI. Together, these findings demonstrate that *HK2* essentiality is shaped by a more complex interplay of intrinsic and extrinsic factors that may in part alter HK1 docking to the OMM, likely clarifying why *HK2* dependence has been masked at least in part over hundreds of CRISPR screens in traditional media.

Nearly all cataloged human cancer lines co-express *HK1* and *HK2*^27^. By contrast, *HK2* shows a more restricted expression profile among normal tissues^25^. This is well illustrated by RNA-Seq data across a broad panel of non-diseased tissue sites from the Adult Genotype Tissue Expression (GTEx) Project (Extended Data Fig. 8a, b)^87^. Given this distinction and the central role of glucose metabolism for cell growth, it is perhaps unsurprising that HK2 has garnered attention as a possible target for cancer therapy^61,88–90^. Non-selective inhibitors such as 2-DG have been limited by toxicity concerns, while efforts to develop more selective HK2 inhibitors have had modest outcomes based on isoform specificity or necessary treatment dosing^44,91–94^. Nonetheless, xenograft studies in mice have offered evidence that *HK2* knockdown (or knockout) can reduce tumor burden for multiple cancer types that also express *HK1*^62,88,95–98^. Beyond direct HK inhibition, there has been interest in developing agents that disrupt HK-VDAC binding as well^61,99,100^. However, our findings suggest that anchoring HK to the OMM or perhaps to other organelles could impair cell growth while eluding HK1/2 specificity issues or the challenge of selectively targeting OMM-detached HK.

### Demand for localized ATP production by glycolysis may underlie the Warburg effect

The question of why aerobic fermentation benefits proliferating cells has received extensive attention, leading to a variety of proposed but debated explanations^20–23^. Since our work indicated that OMM-detached HK correlates with cell growth and also helps facilitate the Warburg effect, we further leveraged conditional *HK2* essentiality to examine several such theories as well. Systematic supplementation with various nutrients, electron acceptors (or donors), and compounds that can either scavenge ROS or impede pyruvate import into mitochondria each failed to restore the relative growth of *HK2*-knockout cells in HPLM. These results suggested that aerobic glycolysis may not be driven by demands to produce biomass, to regenerate NAD^+^, to modulate the NADH-to-NAD^+^ ratio, or to protect against oxidative stress^20,23,72,101^. Metabolite profiling further revealed that loss of *HK2* had little impact on cellular ATP levels regardless of growth in HPLM or RPMI, supporting rationale that proliferating cells are unlikely ATP limited and also challenging the notion that cells exhibit the Warburg effect because ATP synthesis by glycolysis is either more proteome efficient or faster than by respiration^23,84,102^. Moreover, combined rates of ATP synthesis from glycolysis and respiration were unaffected by *HK2* deletion or culture in HPLM versus RPMI, with those from respiration greater in all cases. However, we find that the relative contributions from each pathway vary such that loss of *HK2* severely impairs ATP production rates by glycolysis in HPLM versus RPMI with only a modest impact on those by respiration – an effect that could be rescued by expressing distinct HK1/2 variants based on their respective cytosolic HK activity.

We propose that glycolytic ATP production is compartmentalized to more efficiently support the high energetic demand of at least one essential process that might be in close physical proximity, such as protein synthesis or active ion transport. Although ATP is the only net metabolic output of fermentation, this rationale for the Warburg effect would be overlooked by strictly considering total ATP yield or bulk ATP levels. Furthermore, it does not rely on other theories for aerobic glycolysis attributed to adaptive advantages within a tumor, including immune evasion or a selection pressure (or preparation) for enduring hypoxic environments, which likely act as passenger gains rather than underlying drivers^21,23,84^. The putative role of glycolytic ATP synthesis when oxygen is available may also vary based on cell-dependent demands or the spatial proximity of specific pathways, perhaps reconciling evidence that the Warburg effect is neither a uniformly shared nor unique feature of proliferating cells^103–105^. Notably, prior studies have reported that compartmentalized ATP production – in some cases from glycolysis – can occur in other contexts as well, including skeletal muscle, endothelial cells, neurons, endocrine cells, and worms^106–110^.

### Conditional gene essentiality data can uncover other regulatory nodes of the Warburg effect

Given that HK mediates only the first step of glucose metabolism, we also asked whether our prior screen data could nominate other genes that, like *HK2*, promote the Warburg effect based on CRISPR phenotypes in HPLM versus RPMI^19^. To explore this idea, we first defined a panel of genes spanning all steps of fermentation from glucose uptake to lactate secretion, added others involved in mitochondrial pyruvate metabolism, and then filtered for genes expressed in K562 cells based on reported RNA-seq data^27^. Among the 39 filtered genes, our screen results suggested that nearly 70% were dispensable for cell growth (probability of dependency < 0.5) in HPLM and RPMI, including *HK1* (Extended Data Fig. 9a and Supplementary Table 4). By contrast, 9 genes scored as essential in both conditions: *SLC16A1* and eight that encode glycolytic enzymes *(GPI, ALDOA, TPI, GAPDH, PGK1, PGAM1, ENO1*, and *PKM)* – of which five represented the only expressed isoform in K562 cells for the respective step in the pathway. *SLC16A1* encodes monocarboxylate transporter 1 (MCT1), which has been characterized primarily for proton-coupled transport of lactate and pyruvate across the plasma membrane, though its substrate preference and the direction of net exchange may each vary with context^70,111–113^. In addition, *GPT2* was the only RPMI-essential hit, a conditional phenotype that we previously traced to the unique availability of alanine in HPLM^19^.

We also identified just 3 genes in our panel as selectively essential in HPLM: *HK2*, *PFKP*, and *SLC16A3*. Remarkably, *PFKP* and *SLC16A3* were also each defined as a dependency for less than 2% of the cell lines in DepMap (Extended Data Fig. 9b). Similar to the case for HK2, phosphofructokinase platelet (PFKP) is one of three co-expressed enzymes that can each catalyze the committed step of glycolysis but vary in their biochemical properties and perhaps localization^114^. Moreover, prior work has shown that *PFKP* knockdown can also reduce glucose uptake and lactate secretion in different cancer cell types^115,116^. *SLC16A3* encodes MCT4, which has been proposed to mainly act as a lactate exporter^117^. This role likely supports cellular pH homeostasis and, indeed, MCT4 is highly expressed in tissues that rely on glycolysis for ATP production^66^. Since our results indicated that baseline rates of glucose and lactate exchange were each higher in HPLM versus RPMI, it is interesting to consider the possibility that MCT4 helps to eliminate excess protons from these cells when MCT1-mediated export otherwise becomes saturated.

Notably, a previous report found that *HK2* and *PFKP* were two of just three glycolysis-related genes (of 24 total tested) whose overexpression markedly elevated lactate release from each of two murine cell lines grown in DMEM, while *SLC16A3* expression had a similar effect in one of these lines as well – though other isoforms of HK, PFK1, and MCT were not evaluated^118^. Nonetheless, based on this near total convergence with our screen results despite extensive experimental differences, we propose that aerobic glycolysis is driven by three nodes perhaps most relevant to investigating the Warburg effect: (1) glucose capture by OMM-detached HK, (2) commitment to glycolysis mediated by PFK1, and (3) sufficient removal of net protons generated by glycolysis. Understanding how PFKP mediates a seemingly non-redundant role among the three PFK1 family members may be of interest for future study. This analysis also suggests that CRISPR screening in HPLM versus RPMI could make it possible to determine whether glycolysis-related genes that control aerobic fermentation further depend on natural intrinsic factors in some cases as well.

### Concluding comments

Our study establishes further evidence that conditional essentiality in HPLM can be exploited to gain insights into the molecular basis of biological pathways and how specific proteins contribute to cell growth. Here, this approach uncovers why deletion of *HK2* can differentially impair growth in nutrient conditions more relevant to human physiology, leading to a distinct explanation underlying how aerobic glycolysis benefits proliferating cells. Our findings also suggest additional directions for future work. Targeted *HK2* deletion or CRISPR screening in HPLM across more broadly expanded cell line panels could reveal genomic contributions to *HK2* dependence. Moreover, it is also possible that OMM-detached HK is not diffusely distributed in the bulk cytosol in all contexts, but instead, may be dynamically recruited to subcellular structures that enable compartmentalized ATP synthesis from glycolysis^119^. Elucidating such potential dynamics may reveal candidate pathways that benefit from this localized source of ATP in different cases. Finally, while enforced anchoring or “relocalization” of HK could prove challenging for cancer therapy, it may be possible to apply approaches based on protein thermal stability to identify compounds that preferentially target OMM-detached HK^120,121^ – namely, by screening in *HK2*-knockout cells that express MBD-deficient versus TOM20-fused HK2.

## Materials and Methods

| REAGENT or RESOURCE | SOURCE | IDENTIFIER |
| --- | --- | --- |
| **Antibodies** |  |  |
| Rabbit monoclonal anti-HK1 (C35C4) | Cell Signaling Technology | Cat #: 2024<br>RRID: AB_2116996 |
| Rabbit monoclonal anti-HK2 (C64G5) | Cell Signaling Technology | Cat #: 2867<br>RRID: AB_2232946 |
| Mouse monoclonal anti-GAPDH (0411) | Santa Cruz Biotechnology | Cat #: sc-47724<br>RRID: AB_627678 |
| Mouse monoclonal anti-LAMP2 (H4B4) | Santa Cruz Biotechnology | Cat #: sc-18822<br>RRID: AB_626858 |
| Rabbit monoclonal anti-CALR (D3E6) | Cell Signaling Technology | Cat #: 12238<br>RRID: AB_2688013 |
| Rabbit monoclonal anti-CAT (D4P7B) | Cell Signaling Technology | Cat #: 12980<br>RRID: AB_2798079 |
| Rabbit monoclonal anti-COXIV (3E11) | Cell Signaling Technology | Cat #: 4850<br>RRID: AB_2085424 |
| Rabbit monoclonal anti-GOLGA1 (D8P2K) | Cell Signaling Technology | Cat #: 13192<br>RRID: AB_2798144 |
| Rabbit monoclonal anti-p70 S6K (49D7) | Cell Signaling Technology | Cat #: 2708<br>RRID: AB_390722 |
| Rabbit monoclonal anti-FLAG (D6W5B) | Cell Signaling Technology | Cat #: 14793<br>RRID: AB_2572291 |
| Horse anti-Mouse IgG HRP | Cell Signaling Technology | Cat #: 7076<br>RRID: AB_330924 |
| Goat anti-Rabbit IgG HRP | Cell Signaling Technology | Cat #: 7074<br>RRID: AB_2099233 |
| **Bacterial and Virus Strains** |  |  |
| XL10-Gold Ultracompetent Cells | Agilent | Cat #: 200315 |
| **Chemicals, Peptides, and Recombinant Proteins** |  |  |
| 3X FLAG Peptide | Sigma-Aldrich | Cat #: F4799 |
| Anti-FLAG M2 Affinity Gel | Sigma-Aldrich | Cat #: A2220 |
| Mixture of amino acid standards for metabolomics | Cambridge Isotope Laboratories | Cat #: MSK-A2-1.2 |
| Synthetic medium components | Multiple | See Supplementary Table 1 |
| RPMI 1640, no glucose | Thermo Fisher | Cat #: 11879020 |
| RPMI 1640 | Thermo Fisher | Cat #: 11875093 |
| SILAC RPMI 1640 Flex Media | Thermo Fisher | Cat #: A2494201 |
| RPMI 1640, no glucose, glutamine, vitamin B12, folic acid, phenol red, bicarbonate | US Biological Sciences | Cat #: R9001-02 |
| DMEM, high glucose, GlutaMAX | Thermo Fisher | Cat #: 10566024 |
| DMEM, no glucose | Thermo Fisher | Cat #: 11966025 |
| HPLM | Thermo Fisher | Cat #: A4899101 |
| Penicillin-Streptomycin | Thermo Fisher | Cat #: 15140122 |
| Fetal Bovine Serum (FBS), Heat Inactivated | Thermo Fisher | Cat #: 16140071 |
| X-tremeGENE 9 DNA Transfection Reagent | Sigma-Aldrich | Cat #: 6365779001 |
| Puromycin dihydrochloride | Sigma-Aldrich | Cat #: P7255 |
| Blasticidin (solution) | Invivogen | Cat #: ant-bl-1 |
| Adenosine triphosphate (ATP) | Sigma-Aldrich | Cat #: A0752 |
| Glucose | Thermo Fisher | Cat #: 15023021 |
| Glucose 6-phosphate (G6P) | Sigma-Aldrich | Cat #: G7879 |
| Anti-HA Magnetic Beads | Thermo Fisher | Cat #: 88836 |
| Sodium L-Lactate | Sigma-Aldrich | Cat #: L7022 |
| D-Glucose (U-<sup>13</sup>C<sub>6</sub>, 99%) | Cambridge Isotope Labs | Cat #: CLM-1396 |
| D-Glucose (1,2-<sup>13</sup>C<sub>2</sub>, 99%) | Cambridge Isotope Labs | Cat #: CLM-504 |
| L-Glutamine | Sigma-Aldrich | Cat #: G3126 |
| Sodium Pyruvate | Sigma-Aldrich | Cat #: P2256 |
| L-Alanine | Sigma-Aldrich | Cat #: A7627 |
| N-acetyl-D-glucosamine (GlcNAc) | Sigma-Aldrich | Cat #: A3286 |
| Adenosine | Sigma-Aldrich | Cat #: A9251 |
| Guanosine | Sigma-Aldrich | Cat #: G6264 |
| Hypoxanthine | Sigma-Aldrich | Cat #: H9377 |
| Thymidine | Sigma-Aldrich | Cat #: T1895 |
| Cytidine | Sigma-Aldrich | Cat #: C4654 |
| Uridine | Sigma-Aldrich | Cat #: U3003 |
| Sodium Acetate | Sigma-Aldrich | Cat #: S5636 |
| Dimethyl alpha-ketoglutarate (DM-αKG) | Sigma-Aldrich | Cat #: 349631 |
| Sodium alpha-ketobutyrate (αKB) | Sigma-Aldrich | Cat #: K0875 |
| Sodium 2-hydroxybutyrate (αHB) | Sigma-Aldrich | Cat #: 220116 |
| N-acetyl-L-cysteine (NAC) | Sigma-Aldrich | Cat #: A9165 |
| Duroquinone | Sigma-Aldrich | Cat #: D223204 |
| UK5099 | Selleck Chem | Cat #: S5317 |
| Oligomycin A | Sigma-Aldrich | Cat #: 75351 |
| Rotenone | Sigma-Aldrich | Cat #: R8875 |
| Antimycin A | Sigma-Aldrich | Cat #: A8674 |
| Poly-D-Lysine solution | Sigma-Aldrich | Cat #: A-003-E |
| 2-deoxyglucose (2-DG) | Sigma-Aldrich | Cat #: D8375 |
| Critical Commercial Assays |  |  |
| EndoFree Plasmid Maxi Kit | QIAGEN | Cat #: 12362 |
| Seahorse XFe96 FluxPak Mini | Agilent | Cat #: 103793-100 |
| Deposited Data |  |  |
| Probability of dependency data from genome-wide CRISPR screens in K562 cells | Rossiter et al., 2021<sup>19</sup> | See Supplementary Table 4 |
| Expression data for glycolysis-related gene panel in K562 cells | DepMap, Broad 2024<br>DepMap, 24Q2 Public<sup>27</sup> | See Supplementary Table 4 |
| DepMap gene dependency data for HK isoforms, PFKP, and SLC16A3 | DepMap, Broad 2023<br>DepMap 23Q4 Public<sup>10</sup> | https://depmap.org/portal/data_page/?tab=all<br>Data |
| Expression data for HK isoforms in human cancer cell lines | DepMap, Broad 2024<br>DepMap, 24Q2 Public<sup>27</sup> | https://depmap.org/portal/data_page/?tab=all<br>Data |
| Expression data for HK1 and HK2 in non-disease tissue sites | GTEx Portal<sup>87</sup> | https://www.gtexportal.org/home/downloads/adult-gtex/overview |
| **Experimental Models: Cell Lines** |  |  |
| K562 | J.D. Griffin | ACC-10,<br>RRID_CVCL_0004 |
| NOMO1 | J.D. Griffin | ACC-542,<br>RRID_CVCL_1609 |
| MOLM-13 | CCLE | ACC-554,<br>RRID_CVCL_2119 |
| SUDHL4 | M.A. Shipp | ACC-495,<br>RRID_CVCL_0539 |
| U-937 | D.M. Sabatini | ACC-5,<br>RRID_CVCL_0007 |
| Jurkat | ATCC | ACC-282,<br>RRID_CVCL_0065 |
| Karpas299 | D.M. Sabatini | ACC-31,<br>RRID_CVCL_1324 |
| **Oligonucleotides** |  |  |
| Primers for construction of expression plasmids | This paper | Supplementary Table 2 |
| Individual sgRNA target sequences | This paper | Supplementary Table 2 |
| **Recombinant DNA** |  |  |
| pLJC2-Rap2A-3xFLAG | Cantor et al., 2017<sup>18</sup> | Addgene 87974 |
| pLJC5-Rap2A-3xFLAG | Rossiter et al., 2021<sup>19</sup> | Addgene 163444 |
| pLJC6-Rap2A-3xFLAG | Rossiter et al.,2021<sup>19</sup> | Addgene 163445 |
| pMXs-3xMyc-eGFP-OMP25 | Chen et al., 2016<sup>51</sup> | Addgene 83355 |
| pMXs-3xHA-eGFP-OMP25 | Chen et al., 2016<sup>51</sup> | Addgene 83356 |
| pDONR223-HK1 | Johannessen et al., 2010<sup>123</sup> | Addgene 23730 |
| pDONR223-HK2 | Johannessen et al., 2010<sup>123</sup> | Addgene 23854 |
| pLJC2-HK2-3xFLAG | This paper | Addgene TBD |
| pLJC2-HK2 (D209A, D657A)-3xFLAG | This paper | Addgene TBD |
| pLJC2-TOM20-HK2-3xFLAG | This paper | Addgene TBD |
| pLJC5-HK2-3xFLAG | This paper | Addgene TBD |
| pLJC5-HK1-3xFLAG | This paper | Addgene TBD |
| pLJC5-3xMyc-eGFP-OMP25 | This paper | Addgene TBD |
| pLJC5-3xHA-eGFP-OMP25 | This paper | Addgene TBD |
| pLJC6-EV | Rossiter et al.,2021<sup>19</sup> | Addgene 136454 |
| pLJC6-HK2-3xFLAG | This paper | Addgene TBD |
| pLJC6-HK2 (D209A, D657A)-3xFLAG | This paper | Addgene TBD |
| pLJC6-HK1-3xFLAG | This paper | Addgene TBD |
| pLJC6-HK1 (ΔMBD)-3xFLAG | This paper | Addgene TBD |
| pLJC6-HK2 (ΔMBD)-3xFLAG | This paper | Addgene TBD |
| pLJC6-MBD1-HK2-3xFLAG | This paper | Addgene TBD |
| pLJC6-TOM20-HK2-3xFLAG | This paper | Addgene TBD |
| pLentiCRISPR-v1 | Addgene | Addgene 49535 |
| pLentiCRISPR-v1-sgAAVS1 | Wang et al., 2015<sup>3</sup> | Addgene 70661 |
| pentiCRISPR-v1-sgHK2_14 | This paper | Addgene TBD |
| Software and Algorithms |  |  |
| XCalibur version 4.1 | Thermo Fisher | https://www.thermofisher.com/us/en/home.html |
| R studio version 2023.12.0 | The R Project | https://www.r-project.org/ |
| Prism version 10 | GraphPad | https://www.graphpad.com/ |
| BD FACS Chorus 3.0 | BD Biosciences | https://www.bdbiosciences.com/en-us |
| Wave Controller 2.6 | Agilent | https://www.agilent.com/ |
| Image Studio 5.2 | LI-COR | https://www.licor.com/ |
| Other |  |  |
| SnakeSkin dialysis tubing, 3.5K MWCO | Thermo Fisher | Cat #: PI88244 |
| Z2 Coulter Counter | Beckman | Cat #: 6605700 |
| Multisizer 4e Coulter Counter | Beckman | Cat #: B23005 |
| BD FACSMelody | BD Biosciences | Cat #: NA |
| Seahorse XFe96 Flux Analyzer | Agilent | Cat #: NA |
| LI-COR Odyssey FC | LI-COR | Cat #: 2800 |

### Cell lines

The Jurkat cell line was purchased from ATCC. The following cell lines were kindly provided by: K562 and NOMO1, J. Griffin (Dana Farber Cancer Institute); SUDHL4, M.A. Shipp (Dana Farber Cancer Institute), U-937 and Karpas299, D.M. Sabatini (Institute of Organic Chemistry and Biochemistry of the Czech Academy of Sciences; from lab cell bank); and MOLM-13, Cancer Cell Line Encyclopedia (Broad Institute). Cell lines were verified to be free of mycoplasma contamination and their identities were authenticated by short tandem repeat (STR) profiling.

### Cell culture conditions

The following culture media were used (all contained 0.5% penicillin-streptomycin unless noted):

1. RPMI^+S^: RPMI 1640, no glucose (Thermo Fisher) with 5 mM glucose and 10% FBS.
2. RPMI^+dS^: RPMI 1640, no glucose (Thermo Fisher) with 5 mM glucose and 10% dialyzed FBS.
3. RPMI11^+S^: RPMI1640 (Thermo Fisher) with 10% FBS.
4. RPMI11^+2S^: RPMI1640 (Thermo Fisher) with 20% FBS.
5. RPMI-SH: RPMI 1640 for Seahorse Assays (Supplementary Table 1), no penicillin-streptomycin.
6. DMEM^+S^: DMEM, high glucose, GlutaMAX (Thermo Fisher) with 10% FBS.
7. DMEM^+2S^: DMEM, high glucose, GlutaMAX (Thermo Fisher) with 20% FBS.
8. DMEM5^+S^: DMEM, no glucose (Thermo Fisher) with 5 mM glucose and 10% FBS.
9. HPLM^+dS^: manually prepared HPLM (Supplementary Table 1) with 10% dialyzed FBS.
10. HPLM^+dS^ (Thermo): HPLM (Thermo Fisher) with 10% dialyzed FBS.
11. HPLM-SH: HPLM for Seahorse Assays (Supplementary Table 1), no penicillin-streptomycin.

Using SnakeSkin tubing (Thermo Fisher PI88244), FBS was dialyzed as previously described^18^. Before use, all FBS-supplemented media were sterile-filtered using a bottle-top vacuum filter with cellulose acetate membrane, pore size 0.2 μm (Corning 430626 or Nalgene 290-4520). Cultured cells were maintained at 37°C, atmospheric oxygen, and 5% CO_2_.

### Plasmid construction

All oligonucleotides used in this study are described in Supplementary Table 2.

#### Construction of HK2 knockout plasmid

Sense and antisense oligonucleotides for an *HK2*-targeting single guide RNA (sgRNA) were annealed and then cloned into *BsmBI-*digested pLentiCRISPR-v1.

#### Construction of expression plasmids

The *HK1* gene was amplified from pDONR223-HK1 using primers HK1-F/HK1-R, digested with *PacI-NotI*, and cloned into pLJC5-Rap2A-3xFLAG to generate pLJC5-HK1-3xFLAG. Plasmid pLJC6-HK1-3xFLAG was generated by digesting pLJC5-HK1-3xFLAG with *PacI-NotI* and subcloning into pLJC6-Rap2A-3xFLAG. The HK1(λλMBD)-3xFLAG cDNA was amplified from the pLJC6-HK1-3xFLAG template using primers HK1-MBD/LJC-R, digested with *PacI-NotI,* and then cloned into pLJC6-Rap2A-3xFLAG.

The *HK2* gene was amplified from pDONR223-HK2 using primers HK2-F/HK2-R, digested with *PacI-NotI*, and then cloned into pLJC5-Rap2A-3xFLAG to generate pLJC5-HK2-3xFLAG. The plasmid pLJC6-HK2-3xFLAG contains an sgHK2_14-resistant *HK2* cDNA, which was generated using a 2-step protocol based on overlap extension PCR. In the first step, two fragments were amplified from pLJC5-HK2-3xFLAG using the following primer pairs: LJC-F/HK2_14-R and HK2_14-F/LJC-R. In the second step, the two fragments were pooled in a PCR containing primers LJC-F/LJC-R, then digested with *PacI-NotI*, and cloned into pLJC6-Rap2A-3xFLAG. All other *HK2* expression plasmids were based on this sgHK2_14-resistant *HK2* cDNA.

Plasmid pLJC6-HK2 (D209A, D657A)-3xFLAG was generated by the same two-step overlap extension method described above but instead using the following primer pairs in the first step: LJC-F/HK2-D657A and HK2-D209A/LJC-R. The following cDNAs were amplified, digested with *PacI-NotI*, and then cloned into pLJC6-Rap2A-3xFLAG: (1) HK2(λλMBD)-3xFLAG using primers HK2-MBD/LJC-R; (2) MBD1-HK2-3xFLAG using primers MBD1-HK2/LJC-R; and (3) TOM20-HK2-3xFLAG using primers TOM20-HK2/LJC-R. Plasmids pLJC2-HK2-3xFLAG, pLJC2-HK2 (D209A, D657A)-3xFLAG, and pLJC2-TOM20-HK2-3xFLAG were generated by digesting the pLJC6 plasmids containing each respective cDNA with *PacI-NotI* and then subcloning into pLJC2-Rap2A-3xFLAG.

3xMyc-eGFP-OMP25 and 3xHA-eGFP-OMP25 cDNAs were amplified from the corresponding pMXs plasmid using primers MYC-MITO/MITO-R and HA-MITO/MITO-R, respectively, in turn digested with *PacI-NotI*, and then cloned into pLJC5-Rap2A-3xFLAG.

### Lentivirus production

To produce lentivirus, HEK293T cells in DMEM^+S^ were co-transfected with VSV-G envelope plasmid, Delta-VPR packaging plasmid, and a transfer plasmid (pLJC5, pLJC6, or pLentiCRISPR-v1 backbone) using X-tremeGENE 9 Transfection Reagent (Sigma-Aldrich). The medium was exchanged with fresh DMEM^+2S^ 16 hr after transfection, and the virus-containing supernatant was collected at 48 hr post-transfection, passed through a 0.45 μm filter to eliminate cells, and then stored at −80°C.

### Cell line construction

#### HK2-knockout clonal cell line

To establish an *HK2*-knockout clonal line, K562 cells were seeded at a density of 500,000 cells/mL in a 6-well plate containing 2 mL RPMI11^+S^, 8 μg/mL polybrene, and sgHK2_14-containing lentivirus. Spin infection was carried out by centrifugation at 2,200 RPM for 45 min at 37°C. Following 16-18 hr incubation, cells were pelleted to remove virus and then re-seeded into fresh RPMI11^+S^ for 24 hr. Cells were then pelleted and re-seeded into RPMI11^+S^ containing puromycin for 72 hr. Following selection, the population was single-cell FACS-sorted into 96-well plates containing RPMI11^+2S^ (BD FACSMelody Cell Sorter). After 1.5-2 weeks, cell clones with the desired knockout were identified by immunoblotting. To control for infection, a population of K562 cells was similarly selected following transduction with sgAAVS1-containing lentivirus.

#### HK2-knockout cell lines

The procedure to establish knockout cell lines for short-term growth assays was similar to that used for the clonal cell line except that cells were not FACS-sorted following puromycin selection (5 days post-infection) and with other minor modifications (See **Short-term growth assays**):

1. Cells were transduced in parallel with either sgHK2_14- or sgAAVS1-containing lentivirus.
2. The following cell lines were seeded at a density of 500,000 cells/mL: K562, Karpas299, and MOLM-13.
3. The following cell lines were seeded at a density of 1 million cells/mL: NOMO1, SUDHL4, and U-937.
4. For the HEK293T line, 75,000 cells were seeded in a 6-well plate containing 4 mL DMEM^+S^. Following 24-hr incubation, media were aspirated and replaced with 4 mL DMEM^+S^, 8 μg/mL polybrene, and lentivirus for equivalent spin infection. After 16-18 hr incubation, media were aspirated and replaced with DMEM^+S^ containing puromycin for 72 hr.

The relative depletion of *HK2* in each cell line was assessed by immunoblotting, whereby HK2 signal was normalized by the GAPDH signal in the respective cell lines expressing sgHK2 versus sgAAVS1 (LI-COR Image Studio 5.2).

#### HK cDNA expression cell lines

To establish stable *HK* cDNA expression cell lines, *HK2*-knockout clonal cells were seeded in 6-well plates containing 2 mL of RPMI11^+S^, 8 μg/mL polybrene, and the pLJC6 lentivirus of interest. Spin infection and medium exchange were each performed identically as described above for knockout cell lines. Cells were then pelleted and resuspended into fresh RPMI11^+S^ containing blasticidin for 72 hr. Stable cDNA expression was confirmed by immunoblotting.

#### Mitochondrial immunopurification cell lines

To establish stable expression cell lines for mitochondrial immunopurification, K562 cells (or *HK2*-knockout K562 clonal cells) were seeded at a density of 500,000 cells/mL in 6-well plates containing 2 mL of RPMI11^+S^, 8 μg/mL polybrene, and pLJC5 lentivirus containing either 3xMyc-eGFP-OMP25 or 3xHA-eGFP-OMP25. Spin infection and medium exchange were performed identically as described above for knockout cell lines. Cells were then pelleted and resuspended into fresh RPMI11^+S^ in a 6-well plate. After 48-hr incubation, cells were expanded to 10-cm dishes containing 15 mL of RPM11^+S^ Following 48-hr expansion, cell populations were assessed for GFP signal and then the GFP-positive cells were sorted into RPMI11^+S^ for recovery (BD FACSMelody Cell Sorter).

### Cell lysis for immunoblotting

Cells were centrifuged at 250 *g* for 5 min, resuspended in 1 mL ice-cold PBS, and then centrifuged again at 250 *g* for 5 min at 4°C. Cells were then immediately lysed with ice-cold lysis buffer (40 mM Tris-HCl pH 7.4, 1% Triton X-100, 100 mM NaCl, 5 mM MgCl_2_, 1 tablet of EDTA-free protease inhibitor (Roche 11580800; per 25-mL buffer), and 1 tablet of PhosStop phosphatase inhibitor (Roche 04906845001; per 10-mL buffer)). Cell lysates were cleared by centrifugation at 21,130 *g* for 10 min at 4°C and quantified for protein concentration using an albumin standard (Thermo Fisher 23209) and Bradford reagent (Bio-Rad 5000006). Cell lysate samples were normalized for protein content, denatured upon the addition of 5X sample buffer (Thermo Fisher 39000), resolved by 12% or 4-20% SDS-PAGE, and transferred to a polyvinyl difluoride membrane (Millipore IPVH07850). Membranes were blocked with 5% nonfat dry milk in Tris Buffered Saline with Tween (TBST) for 1 hr at room temperature, and then incubated with primary antibodies in 5% nonfat dry milk in TBST overnight at 4°C. Primary antibodies to the following proteins were used at the indicated dilutions: HK1 (1:1000); HK2 (1:1000);GAPDH (1:1000), LAMP2 (1:500), Calreticulin (CALR) (1:1000), Catalase (CAT) (1:1000), COXIV (1:10,000), GOLGA1 (1:1000), S6K (1:1000), and FLAG (1:1000).

Membranes were washed with TBST three times for 5 min each and then incubated with species-specific HRP-conjugated secondary antibody (1:3000) in 5% nonfat dry milk for 1 hr at room temperature. Membranes were washed again with TBST three times for 5 min each and then visualized with chemiluminescent substrate (Thermo Fisher) on a LI-COR Odyssey FC.

### Short-term growth assays

#### Engineered K562 cell lines

Following at least two passages in RPMI^+S^, cells were pelleted and resuspended to a density of 250,000 cells/mL in either HPLM^+dS^ or RPMI^+dS^. After 48-hr incubation, 3 million cells were pelleted and resuspended to a density of 1 million cells/mL in the respective parent culture medium. From each resuspension, cells were seeded to a final density of 20,000 cells/mL in each of three replicate wells in 6-well plates containing 4 mL of the appropriate culture medium. After 96-hr incubation, cell density measurements were recorded using a Coulter Counter (Beckman Z2 or M4E) with a diameter setting of 8-30 μm.

The short-term growth assay procedure using RPMI- or HPLM-based derivatives was identical to that above with minor modifications:

1. The following were added to RPMI-based media only for the final 96-hr step of the assay with respective stock solutions prepared relative to working concentrations in HPLM: lactate (250X) pyruvate (250X), alanine (500X), and HPLM-specific component pools (see Supplementary Table 1).
2. The following were added to HPLM-based media only for the final 96-hr step of the assay: GlcNAc, adenosine, guanosine, hypoxanthine, thymidine, cytidine, uridine, acetate, DM-

αKG, pyruvate, αKB, αHB, and NAC. Stock solutions of the following individual components were prepared at 100 mM in water: GlcNAc, acetate, DM-αKG, pyruvate, αKB, αHB, and NAC. The stock solution of hypoxanthine was prepared at 10 mM in 0.2 M HCl. Nucleosides were prepared as a single pooled stock solution at 250X (in 0.2 M HCl) relative to the respective working concentrations: adenosine (10 mM), guanosine (10 mM), cytidine (10 mM), uridine (10 mM), and thymidine (1 mM).

1. (3) Modified concentrations of amino acids or salts in HPLM were incorporated at the preceding medium-specific 48-hr passage step. The modified concentration of glutamine in RPMI was also incorporated at the preceding medium-specific 48-hr passage step, with the basal RPMI prepared using SILAC RPMI 1640 Flex Media (Thermo) supplemented with 5 mM glucose, 14 μM phenol red, HPLM-defined glutamine (550 μM), RPMI-defined levels of arginine (1.15 mM) and lysine (218 μM), and 0.5% penicillin-streptomycin (see Supplementary Table 1).

#### K562 cell lines: Drug treatments

The short-term growth assay procedure was identical to that above with minor modifications:

1. Following 1-hr incubation of seeded 6-well plates, compounds were added at specific doses, and then plates were gently shaken for 2 min: Duroquinone (5, 10, or 20 μM), UK5099 (10 μM), or 2-DG (1 mM).
2. For treatment with either duroquinone or UK5099, all wells, including the untreated controls, contained 0.25% DMSO.

Stock solutions of duroquinone and UK5099 were prepared at 40 mM in DMSO. The stock solution of 2-DG was prepared at 400 mM in water.

#### Panel of HK2-knockout cell lines

Cell lines were transduced in parallel with either sgHK2_14- or sgAAVS1-containing lentivirus, and FACS sorting was not performed following puromycin selection. Instead:

1. For the blood cancer cell lines, 3 million cells were pelleted and resuspended to a density of 250,000 cells/mL in 12 mL of RPMI^+S^. After 48-hr incubation (7 days post-infection), 1.75 million cells (x 2) were pelleted and resuspended to a density of ∼300,000 cells/mL in 6 mL of HPLM^+dS^ and RPMI^+dS^. Following 48-hr incubation (9 days post-infection), pools of cells were pelleted and seeded for the 96-hr growth step as described above, with cell density measurements ultimately recorded at 13 days post-infection. Cells grown in HPLM^+dS^ were then lysed for immunoblotting.
2. For the HEK293T line, the cells were detached with 0.05% trypsin-EDTA (Thermo 25300054) and then seeded in duplicate 10-cm culture dishes containing 15 mL DMEM5^+S^. After 48-hr incubation (7 days post-infection), cells from one duplicate culture dish were detached and collected for cell lysis and immunoblotting. Cells from the other duplicate were detached in 2 mL 0.05% trypsin-EDTA and added to 8 mL of fresh DMEM5^+S^. From this resuspension, 1.5 mL was added to a 6-cm culture dish containing 4.5 mL of either HPLM^+dS^ or RPMI^+dS^. After 48-hr incubation (9 days post-infection), cells were detached in 1 mL 0.05% trypsin-EDTA and added to 4 mL of the fresh parent culture medium. From each resuspension, 10,000 cells were seeded for the 96-hr growth step as described above, with cell density measurements ultimately recorded at 13 days post-infection.

### Growth curves

Following at least two passages in RPMI^+S^, cells were pelleted and resuspended to a density of 250,000 cells/mL in either HPLM^+dS^ or RPMI^+dS^. After 48-hr incubation, 1 million cells were pelleted and resuspended to a density of 100,000 cells/mL of the respective parent culture medium. From each resuspension, cells were seeded to a final density of 20,000 cells/mL in each of three replicate T-25 cell culture flasks (Corning 430639) containing 12 mL of the appropriate culture medium. Cell density measurements were recorded every 24 hr using a Coulter Counter (Beckman Z2 or M4E) with a diameter setting of 8-30 μm.

Points for time versus natural log-transformed cell density were plotted in GraphPad Prism and those comprising the exponential (log) phase of the growth curve were fit using linear regression, and the specific growth rate (μ) was defined as the slope of the fit line.

### Metabolite profiling and quantification of metabolite abundance

Liquid chromatography-mass spectrometry (LC-MS) analyses were performed on a QExactive HF benchtop orbitrap mass spectrometer equipped with an Ion Max API source and HESI II probe, coupled to a Vanquish Horizon UHPLC system (Thermo). External mass calibration was performed using positive and negative polarity standard calibration mixtures every 7 days. Acetonitrile was hypergrade for LC-MS (Millipore Sigma), and all other solvents were Optima LC-MS grade (Thermo).

#### Cells: metabolites

At the conclusion of short-term growth assays, a 500 μL aliquot from each well was used to measure cell number and volume via Coulter Counter (Beckman) with a diameter setting of 8-30 μm, and the remaining cells were centrifuged at 250*g* for 5 min, resuspended in 1 mL of ice-cold 0.9% sterile NaCl (Growcells, MSDW1000), and centrifuged at 250 *g* for 5 min at 4°C. Metabolites were extracted in 500 μL of ice-cold 80% methanol containing 500 nM internal amino acid standards (Cambridge Isotope Laboratories). Following a 10-min vortex step and centrifugation at 21,130 *g* for 10 min at 4°C, 2.5 μL from each sample was injected onto a ZIC-pHILIC 2.1 x 150 mm analytical column equipped with a 2.1 x 20 mm guard column (both were 5-μm particle size, Millipore Sigma). Buffer A was 20 mM ammonium carbonate and 40 mM ammonium hydroxide; buffer B was acetonitrile. The chromatographic gradient was run at a flow rate of 0.15 mL/min as follows: 0-20 min: linear gradient from 80 to 20% B; 20.5-28 min: hold at 80% B. The mass spectrometer was operated in full-scan, polarity-switching mode with the spray voltage set to 3.0 kV, the heated capillary held at 275°C, and the HESI probe held at 350°C. The sheath gas flow rate was set to 40 units, the auxiliary gas flow was set to 15 units, and the sweep gas flow was set to 1 unit. The MS data acquisition in positive mode was performed in a range of 50 to 750 mass/charge ratio (m/z), with the resolution set to 120,000, the Automated Gain Control (AGC) target at 10^6^, and the maximum integration time at 20 msec. The settings in negative mode were the same except that the range was instead 70 to 1000 m/z.

For the highly targeted analysis of pyruvate, a tSIM (targeted selected ion monitoring) scan was added with the following settings: resolution set to 120,000, an AGC target of 10^5^, maximum integration time of 200 msec, and isolation window of 1.0 m/z. This tSIM scan was run in negative mode and the target mass was 87.0088.

#### Cells: glucose labeling

Following at least two passages in RPMI^+S^, cells were pelleted and resuspended to a density of 250,000 cells/mL in either HPLM^+dS^ or RPMI^+dS^. After 48-hr incubation, 3 million cells were pelleted and resuspended to a density of 1 million cells/mL of the respective parent culture medium. From each resuspension, cells were seeded to a final density of 20,000 cells/mL in each of three replicate wells in 6-well plates containing 4 mL of appropriate culture medium prepared with 5 mM of either [U-^13^C]-glucose or [1,2-^13^C]-glucose. After 72-hr incubation, metabolite extractions were performed identically to those described above. For the highly targeted analysis of pyruvate isotopologues, tSIM scans were added with the same settings described above and the following target masses: 87.0088 (unlabeled), 88.0121 (M+1-labeled pyruvate), 89.0155 (M+2-labeled pyruvate), and 90.0189 (M+3-labeled pyruvate). To perform natural isotope abundance corrections of isotopologues, the AccuCor code was used^124^.

#### Media: growth curves

At each time point, a 100-μL aliquot from the respective culture flask was collected and centrifuged at 250 *g* for 5 min, and then the resulting supernatant was snap-frozen in liquid nitrogen and stored at −80°C. To extract metabolites from media, samples were thawed on ice and then diluted 1:40 into 50:30:20 methanol:acetonitrile:water containing 500 nM internal amino acid standards (Cambridge Isotope Laboratories). Following a 10-min vortex step and centrifugation at 21,130 *g* for 10 min at 4°C, 2 μL from each sample was injected for analysis as described above for profiling cell samples. For the highly targeted analysis of specific metabolites, tSIM scans were added with the same settings described for cell samples and with the following target masses in negative ionization mode: 87.0088 (pyruvate) and 179.0561 (glucose).

#### HK activity assay evaluation

For the detection of G6P from HK activity assays, reaction mixtures were extracted (See **HK activity assay**), and 5 μL from each sample was injected for analysis as described above for profiling cell samples but using the following chromatographic gradient: 0-10 min: linear gradient from 80 to 20% B; 10-10.5 min: linear gradient from 20 to 80% B; 10.5-17.5 min: hold at 80% B. For the highly targeted analysis of G6P, a tSIM scan was added with the same settings described for cell samples but at a target mass of 259.0224 in negative ionization mode.

#### Identification and quantification

Metabolite identification and quantification were performed with XCalibur v 4.1 (Thermo) using a 10-ppm mass accuracy window and 0.5 min retention time window. To confirm metabolite identities and to enable quantification when desired, a manually constructed library of chemical standards was used. Standards were validated by LC-MS to confirm that they generated robust peaks at the expected m/z ratio, and stock solutions were stored at −80°C. On the day of a given queue, stock solutions were diluted 1:10 in 80% methanol (cell samples) or 50:30:20 methanol:acetonitrile:water (media and HK assay samples) containing 500 nM internal amino acid standards, then vortexed and centrifuged as described for biological samples.

Given that metabolite extraction protocols differed by sample type, the internal standard concentrations in processed samples for metabolites varied: chemical standards (450 nM), media samples (487.5 nM), and cell samples (500 nM). Therefore, the raw peak areas of internal standards within each sample were first normalized to account for these differences. Metabolite quantification was then performed as described elsewhere^18^. For the final concentrations of chemical standards used to quantitate specific metabolites, see Supplementary Table 3.

### Expression and immunoprecipitation of recombinant proteins

For isolation of recombinant proteins, 4 million HEK293T cells were plated in 15-cm culture dishes containing DMEM^+S^. After 24-hr incubation, the cells were transfected with 15 μg of pLJC2 constructs harboring HK2-3xFLAG, HK2 (D209A, D657A)-3xFLAG, or TOM20-HK2-3xFLAG cDNA as described elsewhere^18^. Following an additional 48-hr incubation, cells were rinsed once with ice-cold PBS and then immediately lysed in ice-cold lysis buffer (see **Cell lysis for immunoblotting**). Cell lysates were cleared by centrifugation at 21,130 *g* for 10 min at 4°C. For anti-FLAG immunoprecipitation, FLAG-M2 affinity gel (Sigma-Aldrich) was washed three times in lysis buffer, and then 400 μL of a 50:50 affinity gel slurry was added to a pool of clarified lysates collected from five individual 15-cm culture dishes, and incubated with rotation for 3 hr at 4°C. Following immunoprecipitation, the beads were washed twice in lysis buffer and then four times in lysis buffer containing 500 mM NaCl.

Recombinant protein was then eluted in lysis buffer containing 3x-FLAG peptide (500 μg/mL) for 1 hr with rotation at 4°C. The eluent was isolated by centrifugation at 100 *g* for 4 min at 4°C (Bio-Rad 732-6204), buffer-exchanged (Amicon Ultra 30-kDa MWCO, UFC503024) against 20 volumes of storage buffer (40 mM Tris-HCl pH 7.5, 100 mM NaCl, 2 mM dTT), then mixed with glycerol (final concentration 15% v/v), snap-frozen with liquid nitrogen, and stored at −80°C. Protein samples were quantified using an albumin standard and Bradford reagent. Purified proteins were denatured upon addition of 5X sample buffer and resolved by 12% SDS-PAGE.

### HK activity assay

For all enzyme reactions, the assay buffer was 40 mM Tris-HCl pH 7.4, 5 mM Na_2_HPO_4_, 5 mM MgCl_2_, 2 mM dTT, and 100 μM NaCl. Reactions of recombinant HK2, HK2 (D209A, D657A), and TOM20-HK2 (20-40 nM) with glucose (100 μM) and ATP (100 μM) in assay buffer were carried out at 37°C in a total volume of 100 μL. Following indicated incubation times, a 30 μL aliquot of the reaction was removed and immediately added to 70 μL ice-cold 50:30:20 methanol:acetonitrile:water containing 500 nM internal amino acid standards for metabolite extraction. Samples were then vortexed for 5 min and centrifuged at 21,130 *g* for 1 min at 4°C.

G6P levels generated in each reaction were evaluated by LC-MS analysis of the extracted samples. An identically prepared extraction sample containing only the two HK substrates was used as a background control. G6P levels were quantitated based on a 9-point standard curve using 2-fold dilutions from 20 μM to 0.078125 μM.

### Rapid mitochondrial immunopurification

Following at least two passages in RPMI^+S^, cells expressing 3xHA-eGFP-OMP25 (HA-MITO) or 3xMyc-eGFP-OMP25 (Control-MITO) were pelleted and resuspended to a density of 250,000 cells/mL in either HPLM^+dS^ or RPMI^+dS^. After 48-hr incubation, 7 million cells were pelleted and resuspended to a density of 100,000 cells/mL in 70 mL of the respective parent medium. Following 72-hr incubation, 30 million cells were centrifuged at 250 *g* for 5 min, washed once with 1 mL of ice-cold PBS, and in turn centrifuged at 1000 *g* for 1.5 min at 4°C. Cells were resuspended in 1 mL ice-cold KPBS and then 25 μL of this resuspension was transferred into 90 μL of lysis buffer for extraction of whole-cell lysate (See **Cell lysis for immunoblotting**). Remaining cells were homogenized by one set of 20 strokes in a 2-mL PTFE tissue grinder (VWR 89026-386) and a second set of 10 strokes in a 2-mL dounce tissue grinder (Kimble 885302). After centrifugation at 1000 *g* for 1.5 min at 4°C, ∼800 μL of homogenized supernatant was collected and incubated with 100 μL of anti-HA magnetic beads (Thermo Fisher 88836) pre-washed with KPBS on a head-over-head rotator for 3 min at 4°C. Beads were magnetically isolated (Thermo Fisher DynaMag 12320D) and washed three times with 1 mL ice-cold KPBS. After the final wash step, the beads were lysed in 100 μL of lysis buffer. After magnetically removing the beads, homogenates were cleared by centrifugation at 21,130 *g* for 10 min at 4°C and then denatured upon addition of 5X sample buffer. For immunoblot analyses, 20 μL of whole-cell lysate and 20 μL of corresponding anti-HA eluent were resolved by SDS-PAGE.

Relative mitochondrial enrichment of HK1, HK2, or Flag was assessed by immunoblotting, whereby signals were normalized by the respective COXIV signals in the anti-HA eluent versus in the whole-cell lysate (LI-COR Image Studio 5.2).

### ATP production rate assays

Rates of ATP production from glycolysis and respiration were measured using a Seahorse XFe96 Flux Analyzer (Agilent) according to manufacturer’s protocol for the XF Real-Time ATP Rate Assay Kit with minor modifications. Briefly, XF96 cell culture microplates (Agilent) were coated with 50 μg/mL poly-D-lysine (PDL) and incubated overnight at 37°C in a non-CO_2_ incubator. Microplate wells were then aspirated, rinsed twice with sterile water, and allowed to air-dry for at least 2 hr prior to seeding cells. The Seahorse XFe96 sensor cartridge (Agilent) was hydrated in 200 μL of sterile water and incubated overnight at 37°C in non-CO_2_ incubator.

Following 48-hr incubation in either HPLM^+dS^ or RPMI^+dS^, 150,000 total cells were seeded in replicate wells of PDL-coated microplates containing 50 μL of either HPLM-SH or RPMI-SH (see Supplementary Table 1). Cells were attached by centrifugation at 2,200 RPM for 1 min with no brake and transferred to a non-CO_2_ incubator at 37°C. After 30-min incubation, cells were inspected for attachment before an additional 130 μL of the respective pre-warmed medium was slowly added to each well and then incubated for an additional 15 min at 37°C in a non-CO_2_ incubator. During the cell incubation steps, the sensor cartridge was calibrated with XF Calibrant (Agilent) and ports on the Seahorse Injection Tray were loaded with the following compounds prepared in HPLM-SH or RPMI-SH: port A, 20 μL of oligomycin A (15 μM); port B, 22 μL of rotenone (5 μM) and antimycin A (5 μM). Stock solutions of oligomycin A and rotenone were each prepared at 100 mM in DMSO. The stock solution of antimycin A was prepared at 100 mM in ethanol. For each microplate, a subset of the total wells was not seeded with cells and instead used to calculate the Seahorse XF Buffer Factor of each medium according to manufacturer’s protocol, whereby average buffer factors for HPLM-SH and RPMI-SH were 1.8 and 2.8, respectively.

### Statistical Analysis

*P* values for all comparisons were determined using a two-tailed Welch’s *t*-test. The exact value of *n* and the definition of center and precision measures are provided in the associated figure legends. Bar graphs and most plots were prepared using GraphPad Prism 10; plots comparing *HK1* and *HK2* transcript levels in cancer cell lines, and the probability of dependency for *HK2* in DepMap cell lines were prepared in R. All instances of reported replicates refer to *n* biological samples.

### Net exchange rates

To calculate the rate of net exchange *(q)* for a given metabolite *(M)* between two time points from a respective growth curve, we used the following equation^122^ where μ is the specific growth rate (see **Growth Curves**), *M* is the extracellular metabolite concentration, and *X* is the cell density:

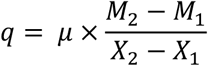

## Acknowledgements

We thank members of the Cantor Lab for upkeep of both the LC-MS system and cell sorter and for helpful discussions, and members of the J. Fan Lab for training and helpful tips on using the Seahorse XFe96 Analyzer. We also thank M. Stefely for organelle figure assets, and K. Tharp and D.M. Sabatini for helpful comments on the manuscript. This work was supported by a grant from the American Cancer Society (RSG-936 21-170-01-TBE to J.R.C). Fellowship support was provided by the NIH (T32HG002760 to K.S.H.), and the University of Wisconsin-Madison Department of Biochemistry to K.S.H., K.M.F., and G.R.C. J.R.C. is a Hartwell Foundation Investigator.

## Author Contributions

K.S.H. and J.R.C. initiated the project and designed the research plan. K.S.H. performed most of the experiments, with assistance from C.M.F., K.M.F., G.R.C., and M.F.M. J.R.C. performed a subset of the growth assays and metabolomics. K.S.H. and J.R.C. analyzed and interpreted the experimental data. J.R.C. wrote the manuscript with assistance from K.S.H. All authors discussed the manuscript.

J.R.C. supervised the studies.

## Competing interests

J.R.C. is an inventor on an issued patent for Human Plasma-Like Medium assigned to the Whitehead Institute (Patent number: US11453858). The remaining authors declare no competing interests.

All data are available in the main text or the supplementary materials. Individual gene knockout and expression plasmids generated in this study will be deposited at Addgene.

Supplementary information is available for this paper (Supplementary Tables 1-4).

## EXTENDED DATA

**Extended Data Fig. 1.**
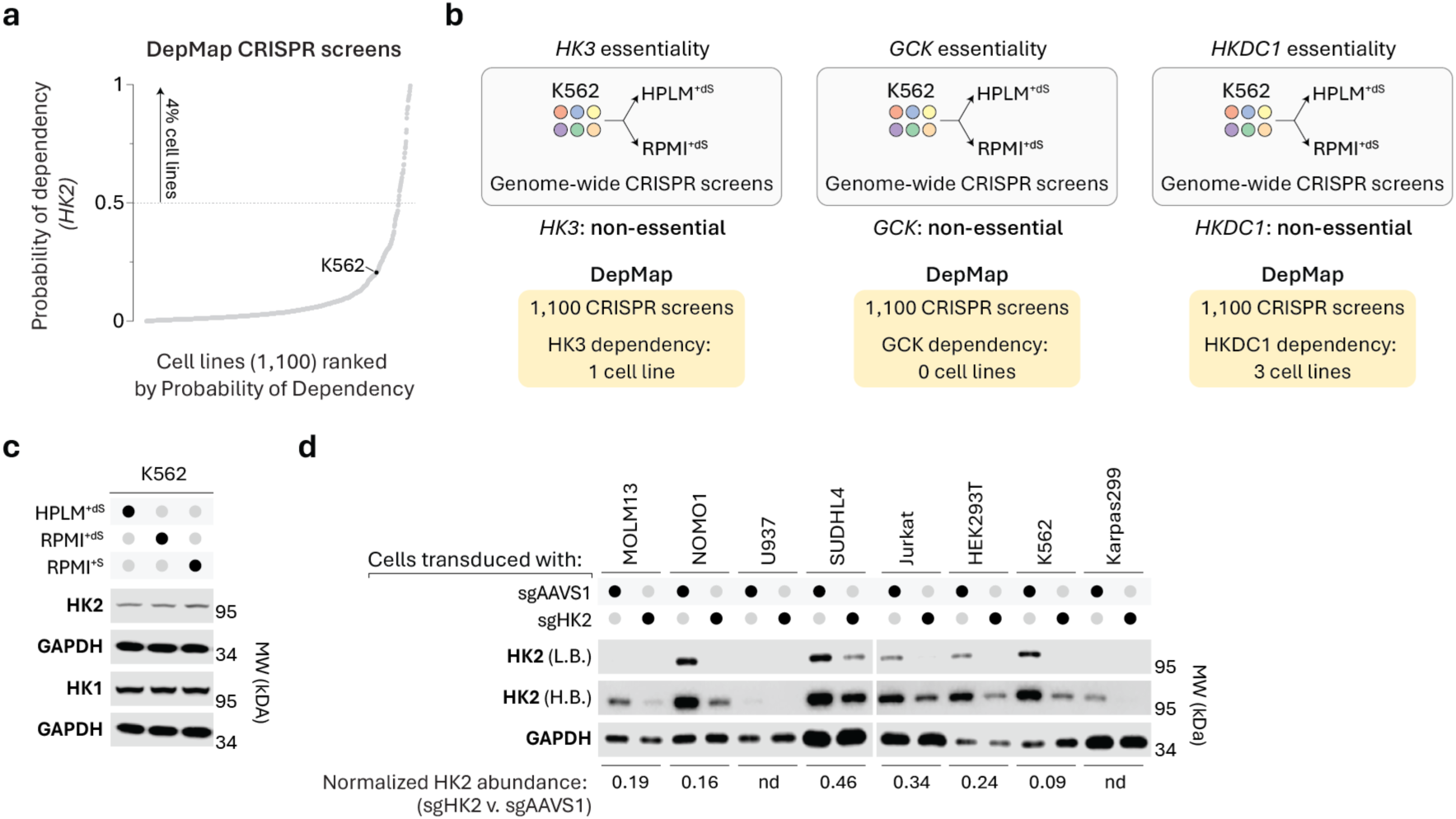
Related to conditional *HK2* essentiality can vary with intrinsic factors. (a) Human cell lines ranked by probability of dependency values for *HK2* across CRISPR screen data cataloged in the DepMap^10^. Probability > 0.5 is the reference threshold for essentiality. (b) Dependency phenotypes for *HK3*, *GCK,* and *HKDC1* from conditional essentiality profiling in K562 cells (top)^19^ and the DepMap (bottom)^10^. (c) Immunoblots for HK1 and HK2. GAPDH served as the loading control in both cases. (d) Immunoblots for expression of HK2 in cell lines transduced with either sgAAVS1 or sgHK2. L.B., low brightness. H.B., high brightness. GAPDH served as the loading control. Bottom row, HK2 signal normalized by GAPDH signal in the respective cell lines expressing sgHK2 versus sgAAVS1 based on immunoblot analysis. nd, not detected.

**Extended Data Fig. 2.**
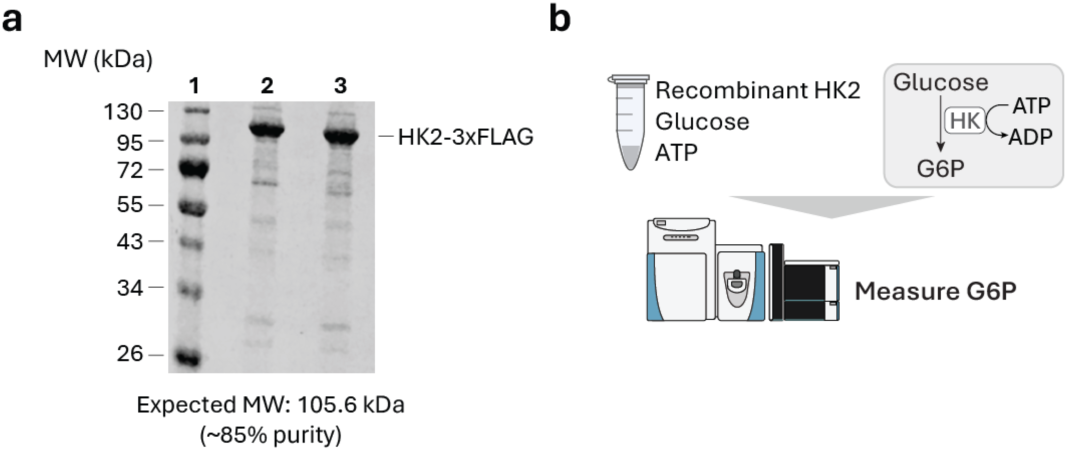
Related to HK2 catalytic activity is necessary to support cell growth in HPLM. (a) Pseudocolor Coomassie-stained gel imaged using a LI-COR Odyssey FC. 1: M.W. standards; 2: wild-type HK2-3xFLAG; 3: HK2 (D209A, D657A)-3xFLAG. (b) Schematic for a method to evaluate HK activity based on measuring glucose 6-phosphate (G6P) production from reactions containing recombinant HK2.

**Extended Data Fig. 3.**
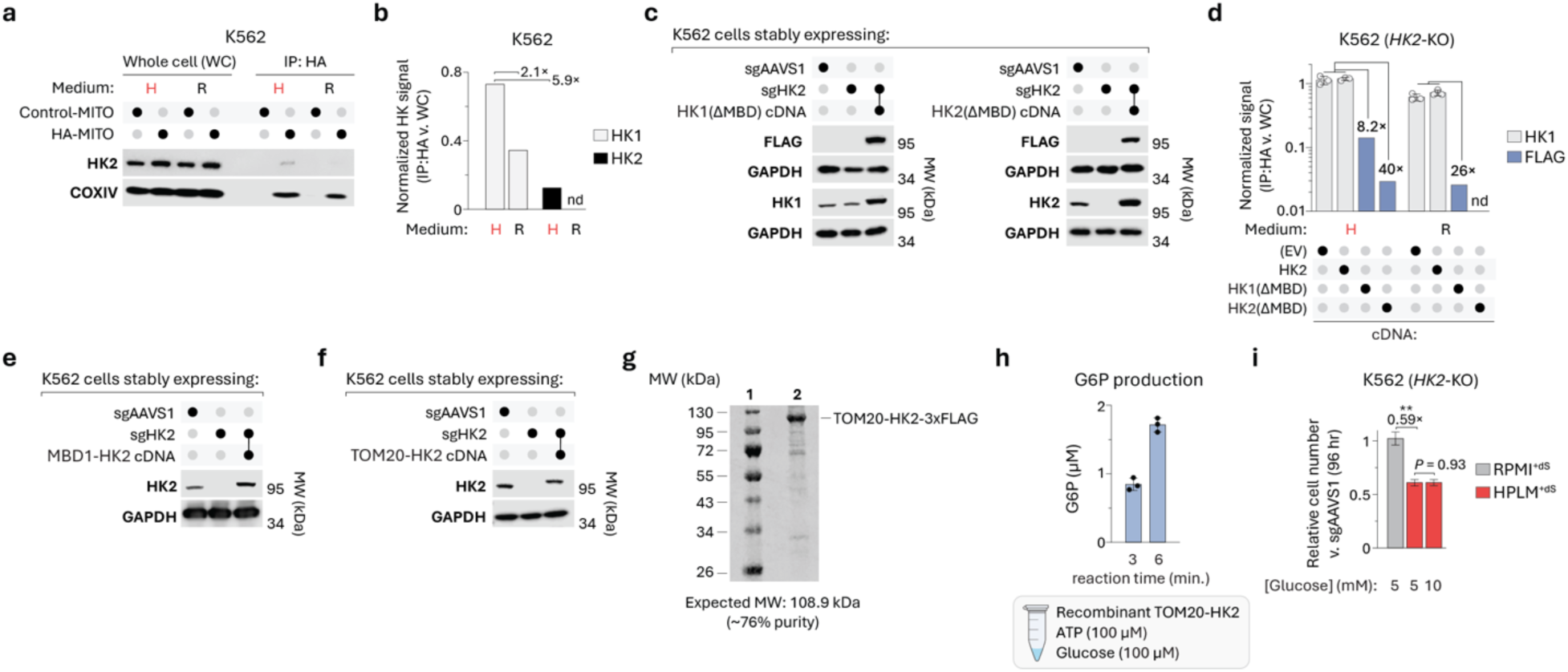
Related to HK1-OMM binding varies with nutrient conditions and HK detachment from mitochondria is necessary to support cell growth. (a) Immunoblots of HK2 and COXIV in whole-cell (WC) lysates and mitochondria immunopurified by HA (IP:HA) from cells expressing either 3xMyc-eGFP-OMP25 (Control-MITO) or 3xHA-eGFP-OMP25 (HA-MITO). COXIV served as a mitochondrial loading control for both HK1 (immunoblot displayed in Fig. 3a) and HK2 (immunoblot displayed here) from the same samples. Immunoblot for HK2 is identical to that displayed in Fig. 3a. (b) HK signal normalized by COXIV signal in IP:HA versus WC lysate based on immunoblot analysis. nd, not detected. (c) Immunoblots for Flag signal and HK1 (left) or Flag signal and HK2 (right). GAPDH served as the loading control in all cases. (d) Gray, HK1 signal normalized by COXIV signal in IP:HA versus WC lysate (mean ± SD, *n* = 3). Blue, Flag signal normalized by COXIV signal in IP:HA versus WC lysate. Values above brackets indicate fold-change between the blue bar and the average of the respective gray bars. (e and f) Immunoblots for HK2. GAPDH served as the loading control. (g) Pseudocolor Coomassie-stained gel imaged using a LI-COR Odyssey FC. 1: M.W. standards; 2: TOM20-HK2-3xFLAG. (h) G6P levels measured from reactions of recombinant TOM20-HK2 with glucose and ATP (mean ± SD, *n* = 3). (i) Relative growth of *HK2*-knockout versus control cells (mean ± SD, *n* = 3, \*\**P* < 0.005).

**Extended Data Fig. 4.**
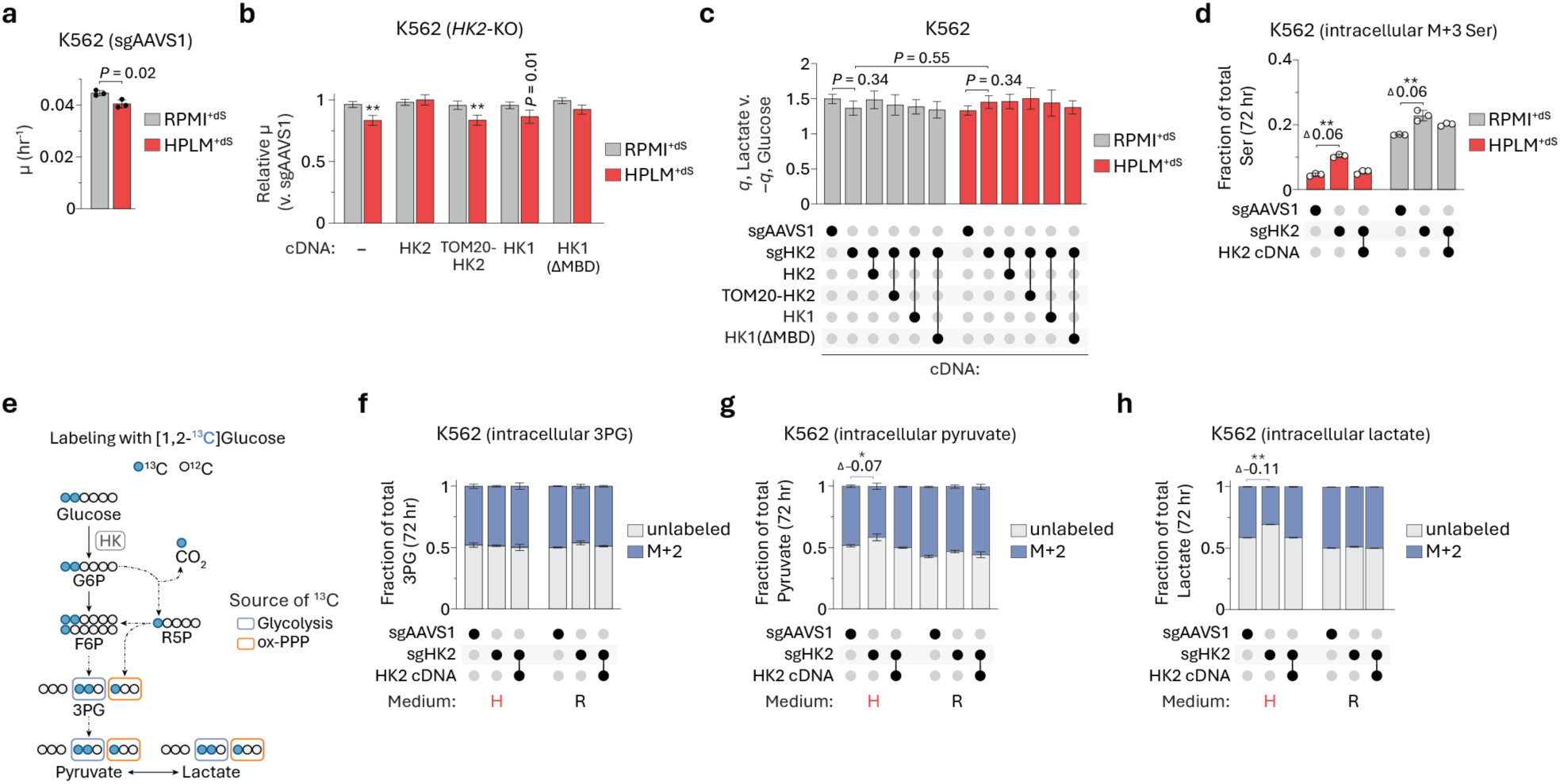
Related to HK detachment from mitochondria supports aerobic glycolysis. (a) Specific growth rates of control cells (mean ± SD, *n* = 3). (b) Relative growth rates of *HK2*-knockout versus control cells (mean ± SD, *n* = 3, \*\**P* < 0.005). (c) Absolute rates of lactate release versus glucose uptake in *HK2*-knockout and control cells (mean ± SEM, *n* = 3). (d) Fractional labeling of serine in *HK2*-knockout and control cells (mean ± SD, *n* = 3, \*\**P* < 0.005). Values above brackets indicate differences in fractional labeling between bars. M+3, incorporation of three 3 ^13^C from [U-^13^C] glucose. (e) Schematic depicting the incorporation of ^13^C from [1,2-^13^C] glucose into various metabolites. Blue box, labeling generated when glucose is metabolized through glycolysis. Orange box, labeling when glucose is metabolized through the oxidative pentose phosphate pathway (ox-PPP). G6P, glucose 6-phosphate. F6P, fructose 6-phosphate. 3PG, 3-phosphoglycerate. (f-h) Fractional labeling of 3PG (f), pyruvate (g), and lactate (h) in *HK2*-knockout and control cells (mean ± SD, *n* = 3, \*\**P* < 0.005, \**P* < 0.05). Values above brackets indicate differences in fractional labeling between bars (g and h). M+2, incorporation of two ^13^C from [1,2-^13^C] glucose.

**Extended Data Fig. 5.**
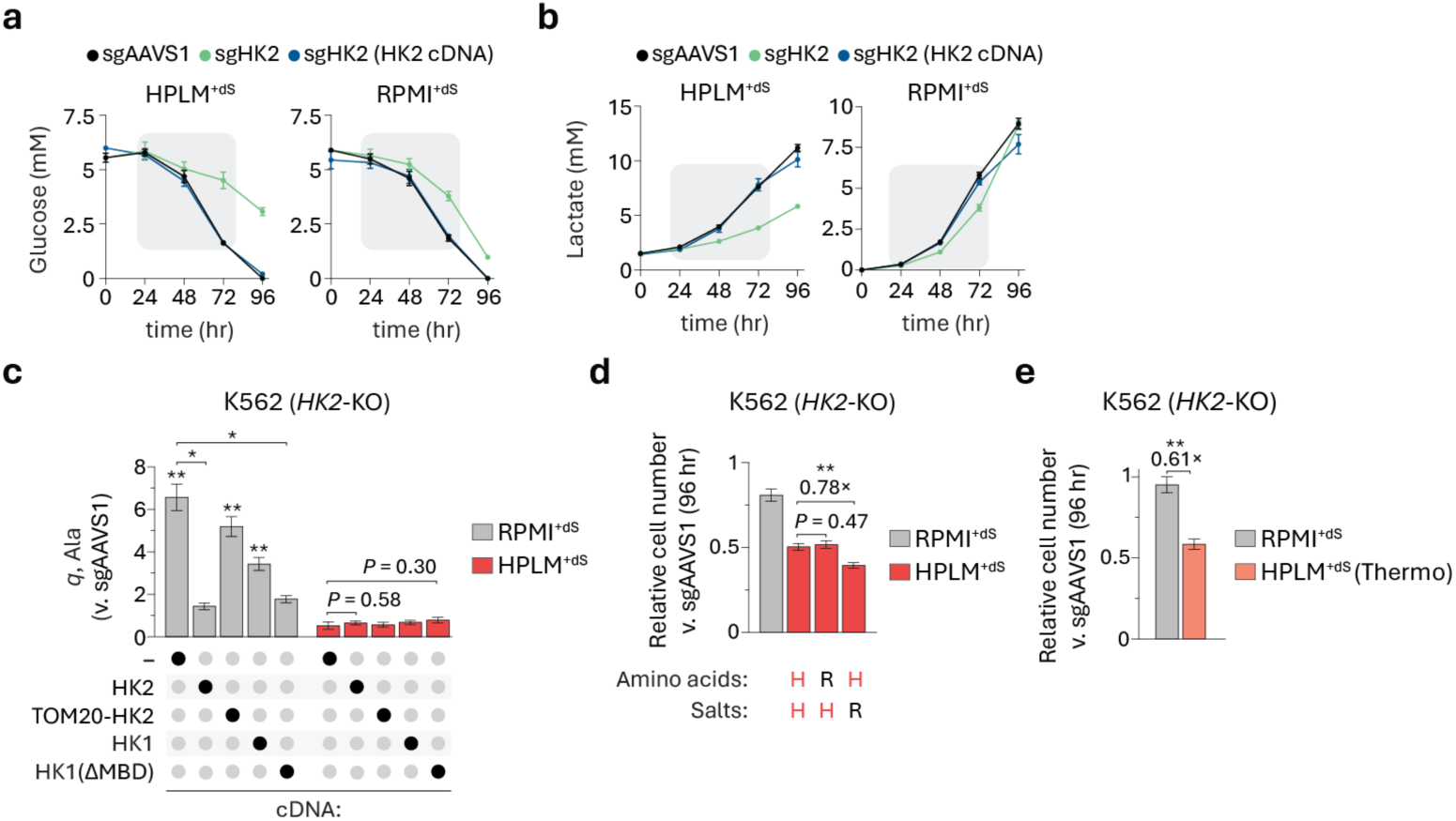
Related to conditional *HK2* essentiality cannot be traced to a direct gene-nutrient interaction. (a and b) Extracellular concentrations of glucose (A) and lactate (B) over the course of growth curves generated for *HK2*-knockout and control cells (mean ± SEM, *n* = 3). Gray box highlights points during log phase. (c) Relative rates of net exchange for alanine in *HK2*-knockout and control cells (mean ± SEM, *n* = 3, \*\**P* < 0.005, \**P* < 0.05). (d and e) Relative growth rates of *HK2*-knockout versus control cells (mean ± SD, *n* = 3, \*\**P* < 0.005). H, HPLM-defined concentrations; R, RPMI-defined concentrations (d). HPLM^+dS^ prepared with HPLM (A4899101, Thermo Fisher Scientific) as the basal medium (e).

**Extended Data Fig. 6.**
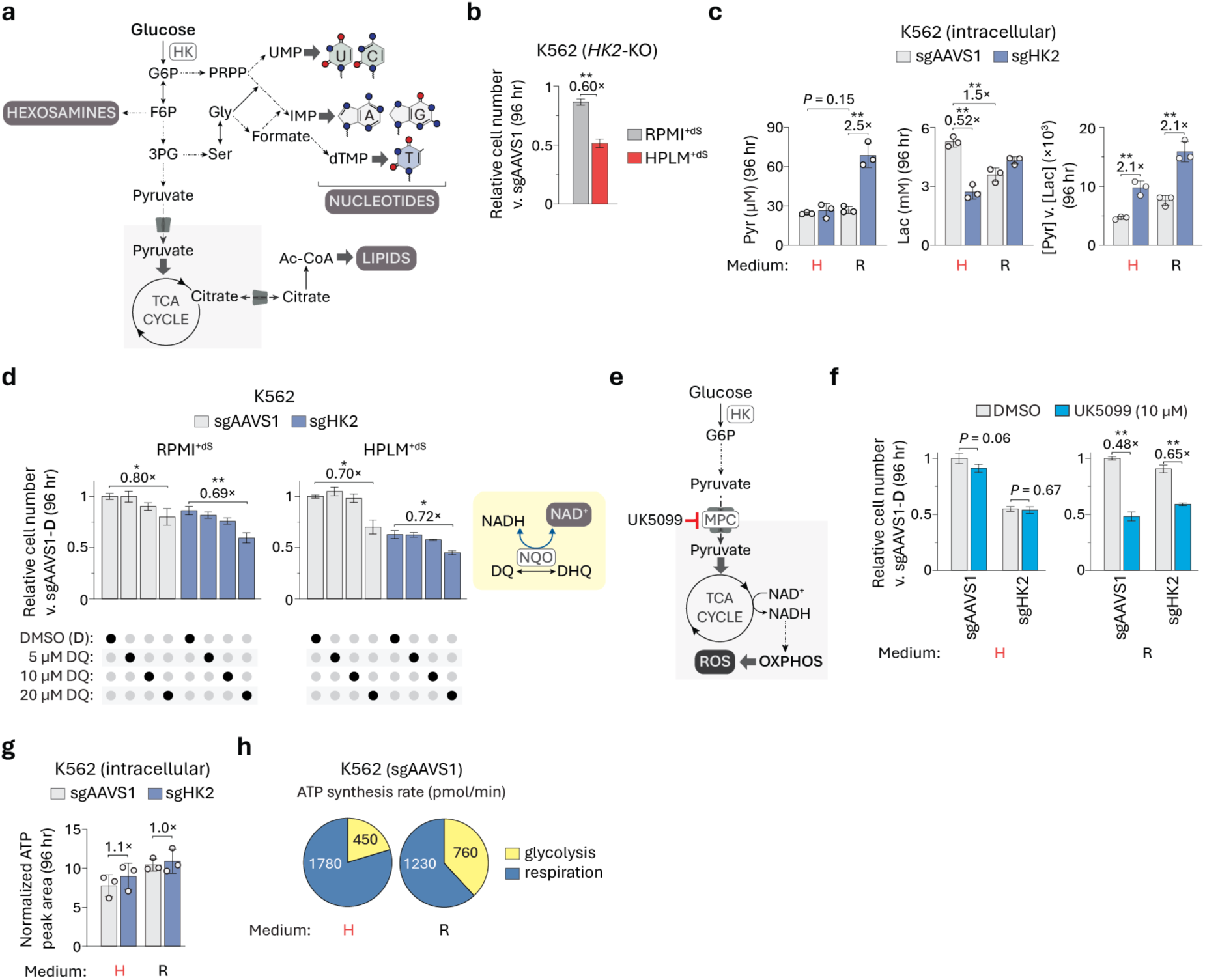
Related to HK detachment from mitochondria drives glycolytic ATP production. (a) Schematic depicting how glucose can be used as a carbon source for the biosynthesis of lipids, nucleotides, and hexosamines. (b) Relative growth rates of *HK2*-knockout versus control cells (mean ± SD, *n* = 3, \*\**P* < 0.005). (c) Cellular concentrations of both pyruvate (left) and lactate (middle) measured at the conclusion of growth assays represented in panel (B). Ratio of cellular pyruvate-to-lactate concentrations (right) (mean ± SD, *n* = 3, \*\**P* < 0.005). (d) Relative growth of *HK2*-knockout and control cells treated with duroquinone (DQ) versus vehicle-treated control cells (mean ± SD, *n* = 3, \*\**P* < 0.005, \**P* < 0.05) (left). DQ as an electron acceptor for the reaction catalyzed by NAD(P)H dehydrogenase [quinone] 1 (NQO). DHQ, durohydroquinone. (e) UK5099 is a small molecule inhibitor of the mitochondrial pyruvate carrier (MPC). (f) Relative growth of *HK2*-knockout and control cells treated with UK5099 versus vehicle-treated control cells (mean ± SD, *n* = 3, \*\**P* < 0.005). (g) Cellular ATP abundances measured at the conclusion of growth assays represented in panel (B) (mean ± SD, *n* = 3). (h) Absolute rates of ATP production by glycolysis (yellow) and by respiration (blue) in control cells. Values depicted are the averages of respective data points (*n* = 6) in Fig. 6i.

**Extended Data Fig. 7.**
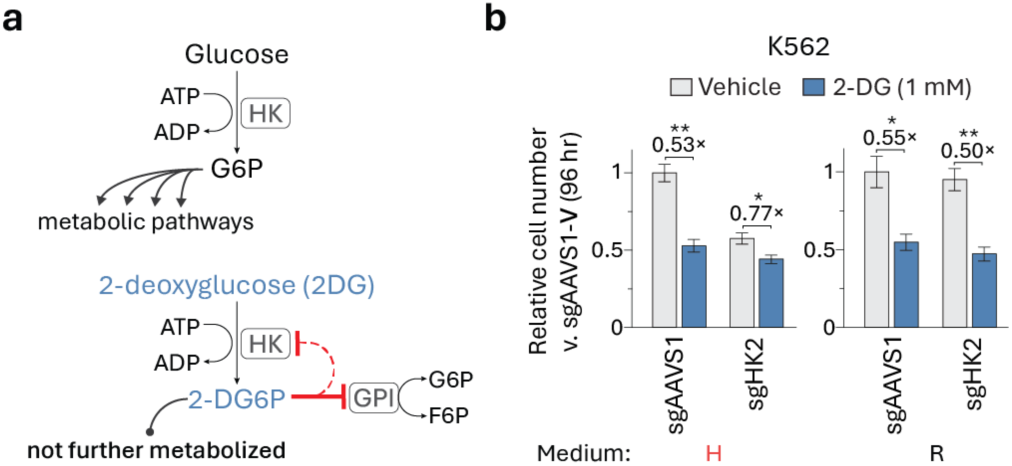
2-deoxyglucose efficacy correlates with HK detachment from mitochondria. (a) 2-deoxyglucose (2-DG) can act as both a competitive and non-competitive inhibitor of HK. 2-DG competes with glucose for HK activity. HK converts 2-DG to 2-deoxyglucose-6-phosphate (2-DG6P), which cannot be further metabolized. 2-DG6P competitively inhibits the glycolytic enzyme glucose-6-phosphate isomerase (GPI) and also mediates allosteric inhibition of HK. (b) Relative growth of *HK2*-knockout and control cells treated with 2-DG versus vehicle-treated control cells (mean ± SD, *n* = 3, \*\**P* < 0.005, \**P* < 0.01).

**Extended Data Fig. 8.**
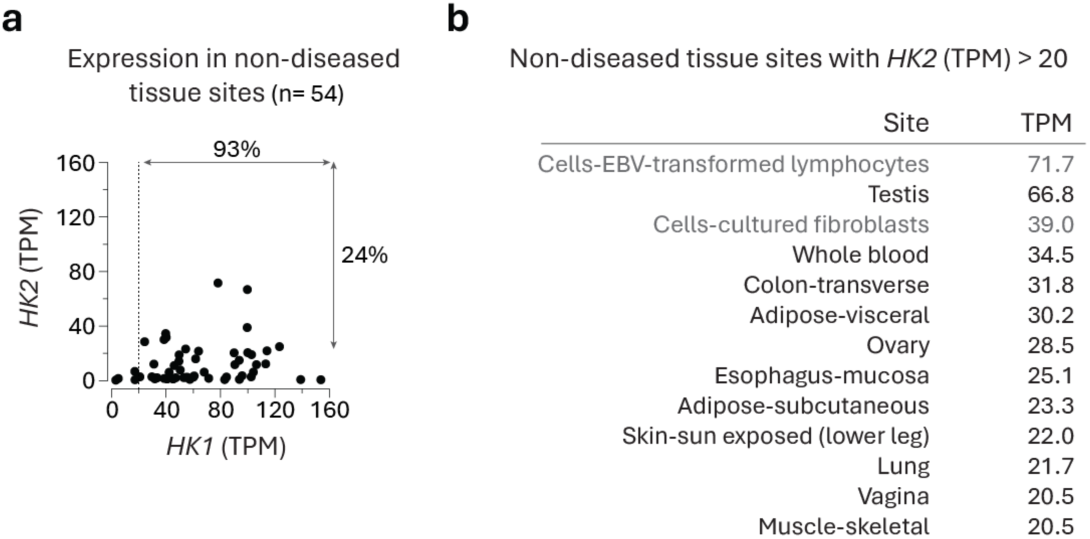
Expression of HK1 and HK2 in non-diseased tissue sites. (a) Comparison between *HK1* and *HK2* RNA transcript levels from the GTEx^87^. TPM, transcripts per million. Dotted lines indicate TPM = 20. (b) Tissue sites with *HK2* TPM greater than 20 (cell lines are in gray text).

**Extended Data Fig. 9.**
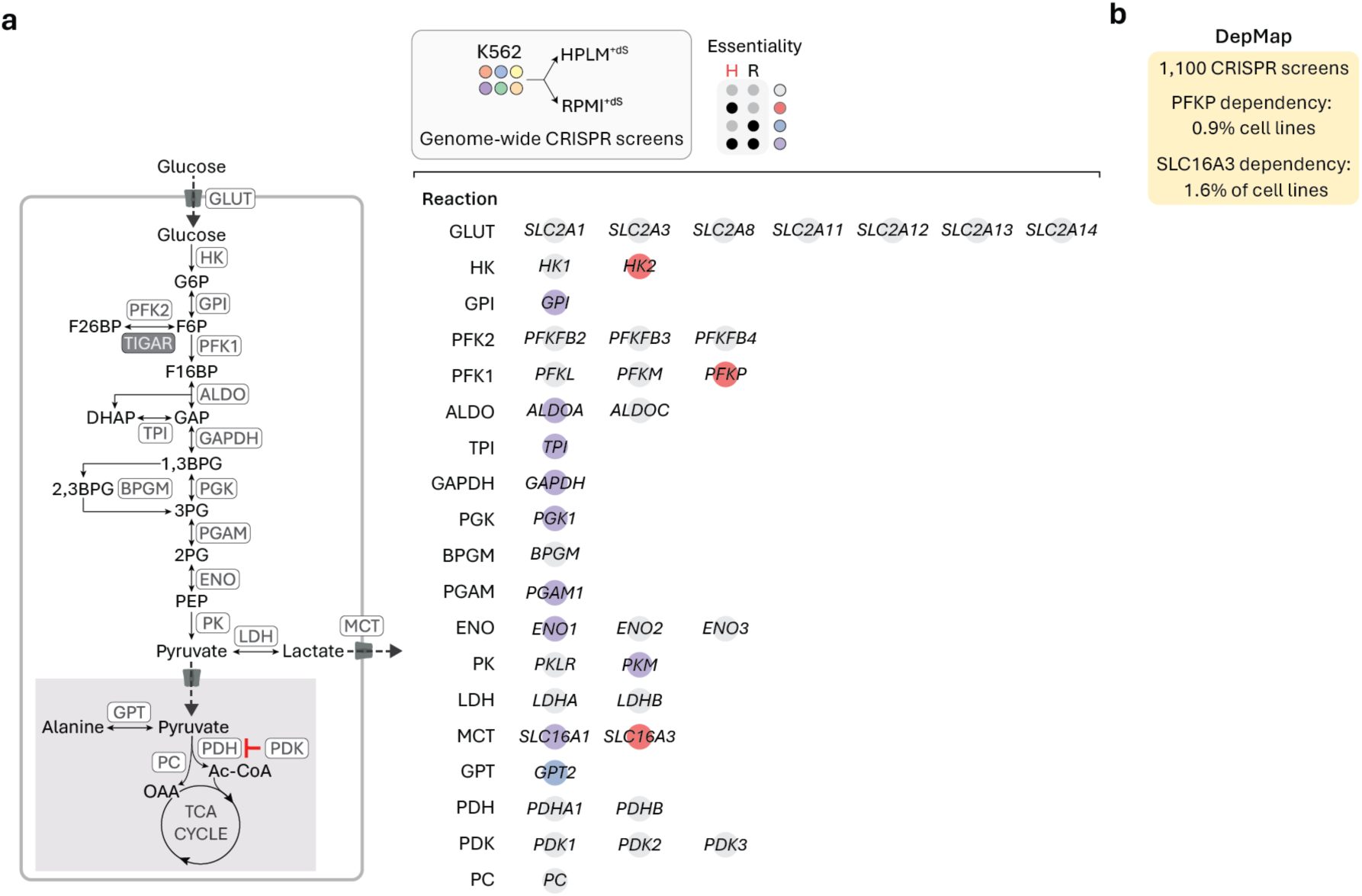
Gene essentiality information from genome-wide CRISPR screens in K562 cells for a panel of glycolysis-related genes. (a) Schematic depicting all steps of fermentation from glucose import to lactate secretion and others that participate in pyruvate metabolism (left). Dependency phenotypes from conditional essentiality profiling in K562 cells^19^ for expressed genes^27^ that can mediate each respective step (right). *TIGAR* is not expressed in K562 cells. Essentiality is defined as probability of dependency > 0.5. (b) Dependency phenotypes for *PFKP* and *SLC16A3* from the DepMap^10^.

## REFERENCES

1. Rancati, G., Moffat, J., Typas, A. & Pavelka, N. Emerging and evolving concepts in gene essentiality. Nat Rev Genet 19, 34–49 (2017).

2. Giaever, G. et al. Functional profiling of the Saccharomyces cerevisiae genome. Nature 418, 387–391 (2002).

3. Wang, T. et al. Identification and characterization of essential genes in the human genome. Science 350, 1096–1101 (2015).

4. Tsherniak, A. et al. Defining a Cancer Dependency Map. Cell 170, 564–576.e16 (2017).

5. Wang, T. et al. Gene Essentiality Profiling Reveals Gene Networks and Synthetic Lethal Interactions with Oncogenic Ras. Cell 168, 890–903.e15 (2017).

6. Wainberg, M. et al. A genome-wide atlas of co-essential modules assigns function to uncharacterized genes. Nat Genet 53, 638–649 (2021).

7. Shi, H., Doench, J. G. & Chi, H. CRISPR screens for functional interrogation of immunity. Nat Rev Immunol 1–18 (2022) doi:10.1038/s41577-022-00802-4.

8. Hart, T. et al. High-Resolution CRISPR Screens Reveal Fitness Genes and Genotype-Specific Cancer Liabilities. Cell 163, 1515–1526 (2015).

9. Hart, T. et al. Evaluation and Design of Genome-Wide CRISPR/SpCas9 Knockout Screens. G3 Genes Genomes Genetics 7, 2719–2727 (2017).

10. Meyers, R. M. et al. Computational correction of copy number effect improves specificity of CRISPR–Cas9 essentiality screens in cancer cells. Nat Genet 49, 1779–1784 (2017).

11. Behan, F. M. et al. Prioritization of cancer therapeutic targets using CRISPR–Cas9 screens. Nature 568, 511–516 (2019).

12. Hillenmeyer, M. E. et al. The Chemical Genomic Portrait of Yeast: Uncovering a Phenotype for All Genes. Science 320, 362–365 (2008).

13. Nichols, R. J. et al. Phenotypic Landscape of a Bacterial Cell. Cell 144, 143–156 (2011).

14. Han, K. et al. CRISPR screens in cancer spheroids identify 3D growth-specific vulnerabilities. Nature 580, 136–141 (2020).

15. Jain, I. H. et al. Genetic Screen for Cell Fitness in High or Low Oxygen Highlights Mitochondrial and Lipid Metabolism. Cell 181, 716–727.e11 (2020).

16. Michl, J. et al. CRISPR-Cas9 screen identifies oxidative phosphorylation as essential for cancer cell survival at low extracellular pH. Cell Reports 38, 110493 (2022).

17. Cantor, J. R. The Rise of Physiologic Media. Trends Cell Biol 29, 854–861 (2019).

18. Cantor, J. R. et al. Physiologic Medium Rewires Cellular Metabolism and Reveals Uric Acid as an Endogenous Inhibitor of UMP Synthase. Cell 169, 258–272.e17 (2017).

19. Rossiter, N. J. et al. CRISPR screens in physiologic medium reveal conditionally essential genes in human cells. Cell Metab 33, 1248–1263.e9 (2021).

20. Vander Heiden, M. G., Cantley, L. C. & Thompson, C. B. Understanding the Warburg Effect: The Metabolic Requirements of Cell Proliferation. Science 324, 1029–1033 (2009).

21. Liberti, M. V. & Locasale, J. W. The Warburg Effect: How Does it Benefit Cancer Cells? Trends Biochem Sci 41, 211–218 (2016).

22. DeBerardinis, R. J. & Chandel, N. S. We need to talk about the Warburg effect. Nat Metabolism 2, 127–129 (2020).

23. 23. Li, Z., Munim, M. B., Sharygin, D. A., Bevis, B. J. & Vander Heiden, M. G. Understanding the Warburg Effect in Cancer. *Cold Spring Harb. Perspect. Med.* Sep 16:a041532 (2024)

24. Fendt, S.-M. 100 years of the Warburg effect: A cancer metabolism endeavor. Cell 187, 3824– 3828 (2024).

25. Wilson, J. E. Isozymes of mammalian hexokinase: structure, subcellular localization and metabolic function. J Exp Biol 206, 2049–2057 (2003).

26. Zapater, J. L., Lednovich, K. R., Khan, Md. W., Pusec, C. M. & Layden, B. T. Hexokinase domain-containing protein-1 in metabolic diseases and beyond. Trends Endocrinol. Metab. 33, 72–84 (2022).

27. Ghandi, M. et al. Next-generation characterization of the Cancer Cell Line Encyclopedia. Nature 569, 503–508 (2019).

28. Ardehali, H. et al. Functional Organization of Mammalian Hexokinase II RETENTION OF CATALYTIC AND REGULATORY FUNCTIONS IN BOTH THE NH2-AND COOH-TERMINAL HALVES . J. Biol. Chem. 271, 1849–1852 (1996).

29. Ferreira, J. C. et al. Linker residues regulate the activity and stability of hexokinase 2, a promising anticancer target. J. Biol. Chem. 296, 100071 (2021).

30. Sui, D. & Wilson, J. E. Structural Determinants for the Intracellular Localization of the Isozymes of Mammalian Hexokinase: Intracellular Localization of Fusion Constructs Incorporating Structural Elements from the Hexokinase Isozymes and the Green Fluorescent Protein. Arch. Biochem. Biophys. 345, 111–125 (1997).

31. Sun, L., Shukair, S., Naik, T. J., Moazed, F. & Ardehali, H. Glucose Phosphorylation and Mitochondrial Binding Are Required for the Protective Effects of Hexokinases I and II▿ †. Mol Cell Biol 28, 1007–1017 (2008).

32. Jin, H. et al. Systematic transcriptional analysis of human cell lines for gene expression landscape and tumor representation. Nat. Commun. 14, 5417 (2023).

33. Pastorino, J. G. & Hoek, J. B. Regulation of hexokinase binding to VDAC. J. Bioenerg. Biomembr. 40, 171–182 (2008).

34. Rose, I. A. & Warms, J. V. B. Mitochondrial Hexokinase RELEASE, REBINDING, AND LOCATION. *J. Biol.* Chem. 242, 1635–1645 (1967).

35. Arora, K. K. & Pedersen, P. L. Functional significance of mitochondrial bound hexokinase in tumor cell metabolism. Evidence for preferential phosphorylation of glucose by intramitochondrially generated ATP. J. Biol. Chem. 263, 17422–17428 (1988).

36. John, S., Weiss, J. N. & Ribalet, B. Subcellular Localization of Hexokinases I and II Directs the Metabolic Fate of Glucose. Plos One 6, e17674 (2011).

37. De Jesus, A. et al. Hexokinase 1 cellular localization regulates the metabolic fate of glucose. Mol Cell 82, 1261–1277.e9 (2022).

38. Pastorino, J. G., Shulga, N. & Hoek, J. B. Mitochondrial Binding of Hexokinase II Inhibits Bax-induced Cytochrome c Release and Apoptosis. J. Biol. Chem. 277, 7610–7618 (2002).

39. Majewski, N. et al. Hexokinase-Mitochondria Interaction Mediated by Akt Is Required to Inhibit Apoptosis in the Presence or Absence of Bax and Bak. Mol. Cell 16, 819–830 (2004).

40. Abu-Hamad, S., Zaid, H., Israelson, A., Nahon, E. & Shoshan-Barmatz, V. Hexokinase-I Protection against Apoptotic Cell Death Is Mediated via Interaction with the Voltage-dependent Anion Channel-1 MAPPING THE SITE OF BINDING. J. Biol. Chem. 283, 13482–13490 (2008).

41. Blaha, C. S. et al. A non-catalytic scaffolding activity of hexokinase 2 contributes to EMT and metastasis. Nat. Commun. 13, 899 (2022).

42. Pilic, J. et al. Hexokinase 1 forms rings that regulate mitochondrial fission during energy stress. Mol. Cell 84, 2732–2746.e5 (2024).

43. Palma, F., Longhi, S., Agostini, D. & Stocchi, V. One-Step Purification of a Fully Active Hexahistidine-Tagged Human Hexokinase Type I Overexpressed in Escherichia coli. Protein Expr. Purif. 22, 38–44 (2001).

44. DeWaal, D. et al. Hexokinase-2 depletion inhibits glycolysis and induces oxidative phosphorylation in hepatocellular carcinoma and sensitizes to metformin. Nat. Commun. 9, 446 (2018).

45. Pastorino, J. G., Hoek, J. B. & Shulga, N. Activation of Glycogen Synthase Kinase 3β Disrupts the Binding of Hexokinase II to Mitochondria by Phosphorylating Voltage-Dependent Anion Channel and Potentiates Chemotherapy-Induced Cytotoxicity. Cancer Res. 65, 10545–10554 (2005).

46. Wang, H. et al. Organization of a functional glycolytic metabolon on mitochondria for metabolic efficiency. Nat. Metab. 6, 1712–1735 (2024).

47. Roberts, D. J., Tan-Sah, V. P., Smith, J. M. & Miyamoto, S. Akt Phosphorylates HK-II at Thr-473 and Increases Mitochondrial HK-II Association to Protect Cardiomyocytes*. J. Biol. Chem. 288, 23798–23806 (2013).

48. Quach, C. H. T. et al. Mild Alkalization Acutely Triggers the Warburg Effect by Enhancing Hexokinase Activity via Voltage-Dependent Anion Channel Binding. PLoS ONE 11, e0159529 (2016).

49. Lafargue, E. et al. Lipid composition of the membrane governs the oligomeric organization of VDAC1. *bioRxiv* 2024.06.26.597124 (2024) doi:10.1101/2024.06.26.597124.

50. Moriya, H. Quantitative nature of overexpression experiments. Mol. Biol. Cell 26, 3932–3939 (2015).

51. Chen, W. W., Freinkman, E., Wang, T., Birsoy, K. & Sabatini, D. M. Absolute Quantification of Matrix Metabolites Reveals the Dynamics of Mitochondrial Metabolism. Cell 166, 1324–1337.e11 (2016).

52. Brito, O. M. de & Scorrano, L. Mitofusin 2 tethers endoplasmic reticulum to mitochondria. Nature 456, 605–610 (2008).

53. Lismont, C., Nordgren, M., Veldhoven, P. P. V. & Fransen, M. Redox interplay between mitochondria and peroxisomes. Front. Cell Dev. Biol. 3, 35 (2015).

54. Chen, W. W., Freinkman, E. & Sabatini, D. M. Rapid immunopurification of mitochondria for metabolite profiling and absolute quantification of matrix metabolites. Nat Protoc 12, 2215–2231 (2017).

55. Schmidt, O., Pfanner, N. & Meisinger, C. Mitochondrial protein import: from proteomics to functional mechanisms. Nat. Rev. Mol. Cell Biol. 11, 655–667 (2010).

56. Kanaji, S., Iwahashi, J., Kida, Y., Sakaguchi, M. & Mihara, K. Characterization of the Signal That Directs Tom20 to the Mitochondrial Outer Membrane. J. Cell Biol. 151, 277–288 (2000).

57. Wu, H. D. et al. Rational design and implementation of a chemically inducible heterotrimerization system. Nat. Methods 17, 928–936 (2020).

58. Koppenol, W. H., Bounds, P. L. & Dang, C. V. Otto Warburg’s contributions to current concepts of cancer metabolism. Nat Rev Cancer 11, 325–337 (2011).

59. Bustamante, E., Morris, H. P. & Pedersen, P. L. Energy metabolism of tumor cells. Requirement for a form of hexokinase with a propensity for mitochondrial binding. J. Biol. Chem. 256, 8699–8704 (1981).

60. Mathupala, S. P., Rempel, A. & Pedersen, P. L. Aberrant Glycolytic Metabolism of Cancer Cells: A Remarkable Coordination of Genetic, Transcriptional, Post-translational, and Mutational Events That Lead to a Critical Role for Type II Hexokinase. J. Bioenerg. Biomembr. 29, 339–343 (1997).

61. Mathupala, S. P., Ko, Y. H. & Pedersen, P. L. Hexokinase-2 bound to mitochondria: Cancer’s stygian link to the “Warburg effect” and a pivotal target for effective therapy. Semin. Cancer Biol. 19, 17–24 (2009).

62. Wolf, A. et al. Hexokinase 2 is a key mediator of aerobic glycolysis and promotes tumor growth in human glioblastoma multiforme. J. Exp. Med. 208, 313–326 (2011).

63. Antoniewicz, M. R. A guide to 13C metabolic flux analysis for the cancer biologist. Exp. Mol. Med. 50, 1–13 (2018).

64. Cantor, J. R. & Sabatini, D. M. Cancer Cell Metabolism: One Hallmark, Many Faces. Cancer Discov 2, 881–898 (2012).

65. Chandel, N. S. Metabolism of Proliferating Cells. Cold Spring Harb. Perspect. Biol. 13, a040618 (2021).

66. Halestrap, A. P. & Wilson, M. C. The monocarboxylate transporter family—Role and regulation. IUBMB Life 64, 109–119 (2012).

67. Parks, S. K., Chiche, J. & Pouysségur, J. Disrupting proton dynamics and energy metabolism for cancer therapy. Nat. Rev. Cancer 13, 611–623 (2013).

68. Rabinowitz, J. D. & Enerbäck, S. Lactate: the ugly duckling of energy metabolism. Nat Metabolism 2, 566–571 (2020).

69. Li, X. et al. Lactate metabolism in human health and disease. Signal Transduct. Target. Ther. 7, 305 (2022).

70. Hong, C. S. et al. MCT1 Modulates Cancer Cell Pyruvate Export and Growth of Tumors that Co-express MCT1 and MCT4. Cell Rep. 14, 1590–1601 (2016).

71. Luengo, A. et al. Increased demand for NAD+ relative to ATP drives aerobic glycolysis. Mol Cell 81, 691–707.e6 (2021).

72. Brand, K. A. & Hermfisse, U. Aerobic glycolysis by proliferating cells: a protective strategy against reactive oxygen species1. FASEB J. 11, 388–395 (1997).

73. Movahed, Z. G., Rastegari-Pouyani, M., Mohammadi, M. & hossein Mansouri, K. Cancer cells change their glucose metabolism to overcome increased ROS: One step from cancer cell to cancer stem cell? Biomed. Pharmacother. 112, 108690 (2019).

74. Schieber, M. & Chandel, N. S. ROS Function in Redox Signaling and Oxidative Stress. Curr. Biol. 24, R453–R462 (2014).

75. Schell, J. C. et al. A Role for the Mitochondrial Pyruvate Carrier as a Repressor of the Warburg Effect and Colon Cancer Cell Growth. Mol. Cell 56, 400–413 (2014).

76. Li, Y. et al. Mitochondrial pyruvate carrier function is negatively linked to Warburg phenotype in vitro and malignant features in esophageal squamous cell carcinomas. Oncotarget 8, 1058–1073 (2016).

77. Yang, C. et al. Glutamine Oxidation Maintains the TCA Cycle and Cell Survival during Impaired Mitochondrial Pyruvate Transport. Mol. Cell 56, 414–424 (2014).

78. Vacanti, N. M. et al. Regulation of Substrate Utilization by the Mitochondrial Pyruvate Carrier. Mol. Cell 56, 425–435 (2014).

79. Molenaar, D., Berlo, R. van, Ridder, D. de & Teusink, B. Shifts in growth strategies reflect tradeoffs in cellular economics. Mol. Syst. Biol. 5, MSB200982 (2009).

80. Vazquez, A., Liu, J., Zhou, Y. & Oltvai, Z. N. Catabolic efficiency of aerobic glycolysis: The Warburg effect revisited. BMC Syst. Biol. 4, 58 (2010).

81. Basan, M. et al. Overflow metabolism in Escherichia coli results from efficient proteome allocation. Nature 528, 99–104 (2015).

82. Wortel, M. T., Noor, E., Ferris, M., Bruggeman, F. J. & Liebermeister, W. Metabolic enzyme cost explains variable trade-offs between microbial growth rate and yield. PLoS Comput. Biol. 14, e1006010 (2018).

83. Kukurugya, M. A., Rosset, S. & Titov, D. V. The Warburg Effect is the result of faster ATP production by glycolysis than respiration. Proc. Natl. Acad. Sci. 121, e2409509121 (2024).

84. Shen, Y. et al. Mitochondrial ATP generation is more proteome efficient than glycolysis. Nat. Chem. Biol. 1–10 (2024) doi:10.1038/s41589-024-01571-y.

85. Vander Heiden, M. G. Targeting cancer metabolism: a therapeutic window opens. Nat Rev Drug Discov 10, 671–684 (2011).

86. Pajak, B. et al. 2-Deoxy-d-Glucose and Its Analogs: From Diagnostic to Therapeutic Agents. Int. J. Mol. Sci. 21, 234 (2019).

87. Lonsdale, J. et al. The Genotype-Tissue Expression (GTEx) project. Nat Genet 45, 580–585 (2013).

88. Patra, K. C. et al. Hexokinase 2 Is Required for Tumor Initiation and Maintenance and Its Systemic Deletion Is Therapeutic in Mouse Models of Cancer. Cancer Cell 24, 213–228 (2013).

89. Galluzzi, L., Kepp, O., Vander Heiden, M. G. & Kroemer, G. Metabolic targets for cancer therapy. Nat. Rev. Drug Discov. 12, 829–846 (2013).

90. Shan, W., Zhou, Y. & Tam, K. Y. The development of small-molecule inhibitors targeting hexokinase 2. Drug Discov. Today 27, 2574–2585 (2022).

91. Lin, H. et al. Discovery of a Novel 2,6-Disubstituted Glucosamine Series of Potent and Selective Hexokinase 2 Inhibitors. Acs Med Chem Lett 7, 217–222 (2016).

92. Li, W. et al. Benserazide, a dopadecarboxylase inhibitor, suppresses tumor growth by targeting hexokinase 2. J. Exp. Clin. Cancer Res. 36, 58 (2017).

93. Liu, Y. et al. Structure based discovery of novel hexokinase 2 inhibitors. Bioorg. Chem. 96, 103609 (2020).

94. Zheng, M. et al. Novel selective hexokinase 2 inhibitor Benitrobenrazide blocks cancer cells growth by targeting glycolysis. Pharmacol. Res. 164, 105367 (2021).

95. Wang, L. et al. Hexokinase 2-Mediated Warburg Effect Is Required for PTEN- and p53-Deficiency-Driven Prostate Cancer Growth. Cell Rep. 8, 1461–1474 (2014).

96. Liu, H. et al. Hexokinase 2 (HK2), the tumor promoter in glioma, is downregulated by miR-218/Bmi1 pathway. PLoS ONE 12, e0189353 (2017).

97. Anderson, M., Marayati, R., Moffitt, R. & Yeh, J. J. Hexokinase 2 promotes tumor growth and metastasis by regulating lactate production in pancreatic cancer. Oncotarget 8, 56081–56094 (2016).

98. Liu, W., Li, W., Liu, H. & Yu, X. Xanthohumol inhibits colorectal cancer cells via downregulation of Hexokinases II-mediated glycolysis. Int. J. Biol. Sci. 15, 2497–2508 (2019).

99. Shoshan-Barmatz, V. & Mizrachi, D. VDAC1: from structure to cancer therapy. Front. Oncol. 2, 164 (2012).

100. Krasnov, G. S., Dmitriev, A. A., Lakunina, V. A., Kirpiy, A. A. & Kudryavtseva, A. V. Targeting VDAC-bound hexokinase II: a promising approach for concomitant anti-cancer therapy. Expert Opin. Ther. Targets 17, 1221–1233 (2013).

101. Wang, Y. et al. Saturation of the mitochondrial NADH shuttles drives aerobic glycolysis in proliferating cells. Mol. Cell 82, 3270–3283.e9 (2022).

102. Vander Heiden, M. G. & DeBerardinis, R. J. Understanding the Intersections between Metabolism and Cancer Biology. Cell 168, 657–669 (2017).

103. Chen, P.-H. et al. Metabolic Diversity in Human Non-Small Cell Lung Cancer Cells. Mol. Cell 76, 838–851.e5 (2019).

104. Hardie, D. G. 100 years of the Warburg effect: a historical perspective. *Endocr.-Relat*. Cancer 29, T1–T13 (2022).

105. Viegas, F. O. & Neuhauss, S. C. F. A Metabolic Landscape for Maintaining Retina Integrity and Function. Front. Mol. Neurosci. 14, 656000 (2021).

106. Han, J. W., Thieleczek, R., Varsanyi, M. & Heilmeyer, L. M. G. Compartmentalized ATP synthesis in skeletal muscle triads. Biochemistry 31, 377–384 (1992).

107. Zala, D. et al. Vesicular Glycolysis Provides On-Board Energy for Fast Axonal Transport. Cell 152, 479–491 (2013).

108. De Bock, K. et al. Role of PFKFB3-Driven Glycolysis in Vessel Sprouting. Cell 154, 651–663 (2013).

109. Kelley, L. C. et al. Adaptive F-Actin Polymerization and Localized ATP Production Drive Basement Membrane Invasion in the Absence of MMPs. Dev. Cell 48, 313–328.e8 (2019).

110. Ho, T., Potapenko, E., Davis, D. B. & Merrins, M. J. A plasma membrane-associated glycolytic metabolon is functionally coupled to KATP channels in pancreatic α and β cells from humans and mice. Cell Rep. 42, 112394 (2023).

111. Doherty, J. R. et al. Blocking Lactate Export by Inhibiting the Myc Target MCT1 Disables Glycolysis and Glutathione Synthesis. Cancer Res. 74, 908–920 (2014).

112. Faubert, B. et al. Lactate Metabolism in Human Lung Tumors. Cell 171, 358–371.e9 (2017).

113. Bosshart, P. D., Charles, R.-P., Garibsingh, R.-A. A., Schlessinger, A. & Fotiadis, D. SLC16 Family: From Atomic Structure to Human Disease. Trends Biochem. Sci. 46, 28–40 (2021).

114. Campos, M. & Albrecht, L. V. Hitting the Sweet Spot: How Glucose Metabolism Is Orchestrated in Space and Time by Phosphofructokinase-1. Cancers 16, 16 (2023).

115. Lee, J.-H. et al. Stabilization of phosphofructokinase 1 platelet isoform by AKT promotes tumorigenesis. Nat. Commun. 8, 949 (2017).

116. Huang, Y. et al. p53-responsive CMBL reprograms glucose metabolism and suppresses cancer development by destabilizing phosphofructokinase PFKP. Cell Rep. 42, 113426 (2023).

117. Contreras-Baeza, Y. et al. Monocarboxylate transporter 4 (MCT4) is a high affinity transporter capable of exporting lactate in high-lactate microenvironments. J. Biol. Chem. 294, 20135–20147 (2019).

118. Tanner, L. B. et al. Four Key Steps Control Glycolytic Flux in Mammalian Cells. Cell Syst 7, 49–62.e8 (2018).

119. Fuller, G. G. & Kim, J. K. Compartmentalization and metabolic regulation of glycolysis. J. Cell Sci. 134, (2021).

120. Gaetani, M. et al. Proteome Integral Solubility Alteration: A High-Throughput Proteomics Assay for Target Deconvolution. J. Proteome Res. 18, 4027–4037 (2019).

121. 121. Vranken, J. G. V. et al. Large-scale characterization of drug mechanism of action using proteome-wide thermal shift assays. *bioRxiv* 2024.01.26.577428 (2024) doi:10.1101/2024.01.26.577428.

122. Shuler, M. L., Mufti, N., Donaldson, M. & Taticek, R. A bioreactor experiment for the senior laboratory. Chemical Engineering Education (1994).

123. Johannessen, C. M. et al. COT drives resistance to RAF inhibition through MAP kinase pathway reactivation. Nature 468, 968–972 (2010).

124. Su, X., Lu, W. & Rabinowitz, J. D. Metabolite Spectral Accuracy on Orbitraps. Anal. Chem. 89, 5940–5948 (2017).

